# Hebb’s Vision: The Structural Underpinnings of Hebbian Assemblies

**DOI:** 10.1101/2025.04.24.649900

**Authors:** J Wagner-Carena, S Kate, T Riordan, R Abbasi-Asl, J Aman, A Amster, AL Bodor, D Brittain, JA Buchanan, MA Buice, DJ Bumbarger, F Collman, NM da Costa, DJ Denman, SEJ de Vries, E Joyce, D Kapner, CW King, JD Larkin, J Lecoq, G Mahalingam, D Millman, J Mölter, C Morrison, RC Reid, CM Schneider-Mizell, S Daniel, S Suckow, KT Takasaki, M Takeno, R Torres, D Vumbaco, J Waters, DG Wyrick, W Yin, J Zhuang, S Mihalas, S Berteau

**Affiliations:** Allen Institute, Seattle, WA, USA; Department of Molecular and Cellular Biology, Harvard University, Cambridge, MA, USA; Neuroscience Graduate Program, University of California, San Francisco, San Francisco, CA, USA; Department of Neurology, University of California, San Francisco, San Francisco, CA, USA; Department of Biomedical Engineering, Georgia Institute of Technology, Atlanta, GA, USA; Department of Mathematics, School of Computation, Information and Technology, Technical University of Munich, Germany

## Abstract

In 1949, Donald Hebb proposed that groups of neurons that activate stereotypically form the organizational building blocks of perception, cognition, and behavior. Finding the structural underpinning of such assemblies has been technically challenging, due to a lack of large-scale structure-activity maps. Here, we analyze this relation using a novel dataset that links in vivo optical physiology to connectivity using postmortem electron microscopy (EM). From the fluorescence traces, we extract neural assemblies from higher-order correlations in neural activity. Physiologically, we show that these assemblies exhibit properties consistent with Hebb’s theory, including more reliable responses to repeated natural movie inputs than size-matched random ensembles and superior decoding of visual stimuli. Structurally, we find that neurons that participate in assemblies are significantly more integrated into the structural network than those that do not. Contrary to Hebb’s original prediction, we do not observe a marked increase in the strength of monosynaptic excitatory connections between cells participating in the same assembly. However, we find significantly stronger indirect feed-forward inhibitory connections targeting cells in other assemblies. These results show that assemblies can be useful components of perception, and, surprisingly, they are delineated by mutual inhibition.

## 2 Introduction

Since Hebb’s 1949 monograph, The Organization of Behavior[1], cell assemblies have retained a persistent place in the imagination of the neuroscience community, both as a prospective unit of functional organization and as a compellingly likely consequence of simple rules of synaptic plasticity, such as the “fire-together wire-together” synapses dubbed ‘Hebbian Synapses’ by Yves Frégnac in 1986 [2]. Throughout this work, we follow Hebb in using the term assembly to refer to overlapping sets of neurons that activate in a reproducible pattern with high fidelity; in much of the contemporary literature, such groups are also called cell ensembles or simply ensembles. The theory continues to stimulate research on the activity-related aspects of assemblies, and the organization of these processing modules is thought to operate through stable recurrent activity [3, 4]. Incorporating a behavioral approach, experimental evidence implies that the response of discrete cell populations akin to assemblies may also have a causal link with motor functions [5]. In particular, there has been confirmation of sequential activation within neural assemblies of the primary visual cortex (V1) [6] as well as ongoing ‘replay’ of coactivity in the absence of stimuli [7]. There has also been some limited evidence of cell assemblies’ stability over long periods of time [8].

However, research examining the structural aspects of Hebbian cell assemblies has primarily focused on the potentiation of synapses between excitatory neurons. This emphasis is perhaps not surprising, given that this potentiation has a place of prominence in Hebb’s original formulation of the theory [1]. In 1963 [9], Hebb himself acknowledged that his excitation-only formulation of assembly theory was a concession to the state of research on inhibitory synapses at that point in time; synaptic inhibition of neural activity was not confirmed until 1952 [10]. Over the following decades, the association between Hebbian assemblies and excitatory potentiation has itself been reinforced by the computational plausibility of excitatory plasticity as a mechanism of assembly formation [11, 12, 13], and the discovery of Hebbian synapses. However, such changes were never a necessary precondition for assembly formation. Mechanistically, there exists a broad range of plausible solutions to assembly formation, ranging from excitatory modulation against a backdrop of relatively stable inhibitory strengths to the opposite, in which formation relies solely on inhibitory modulation contrasting with stable excitatory connections [14].

The approach of this work, using large-scale recordings of individual neurons to identify and analyze assembly function, enters a developing tradition in the literature of population dynamics [15, 16, 17, 18, 19, 20]. However, due to limitations in structural analysis, few studies can relate the structure of assemblies to their function, as reviewed by [14]. Electrophysiological datasets can allow for a highly reliable inference of connectivity in the case of multi-patch recordings, but with very small numbers of cells in any given study. Extracellular recordings can overcome this limitation, but produce biased connectivity estimates, and therefore cannot be considered a gold standard.

To relate the correlated activity of a large number of neurons to their connections, we used a novel large-scale multi-modal dataset: the Allen Brain Observatory V1 Deep Dive (V1DD) (Fig. 1A). V1DD offers a combination of Ca^2+^ fluorescent recordings and detailed EM reconstruction of neurons and synapses, including post-synaptic density volumes. Taking advantage of advances in optical imaging techniques [21], V1DD provides multiple scans of high-quality simultaneous two-photon calcium imaging (2PCI) recordings of thousands of excitatory neurons within the mouse primary visual cortex (Fig. 1B). In addition to its functional recordings, V1DD also contains electron microscopy (EM) of the same tissue volume, which has uncovered the fine-scale anatomy of the cubic millimeter volume of the brain (Fig. 1C). EM has been employed extensively in large-scale datasets to map drosophila [22, 23], worm [24], mouse [25, 26], and even the human brain [27]. The combination of these two imaging modalities has been applied in very few datasets [28, 29, 26], particularly at this scale.

**Figure 1.**
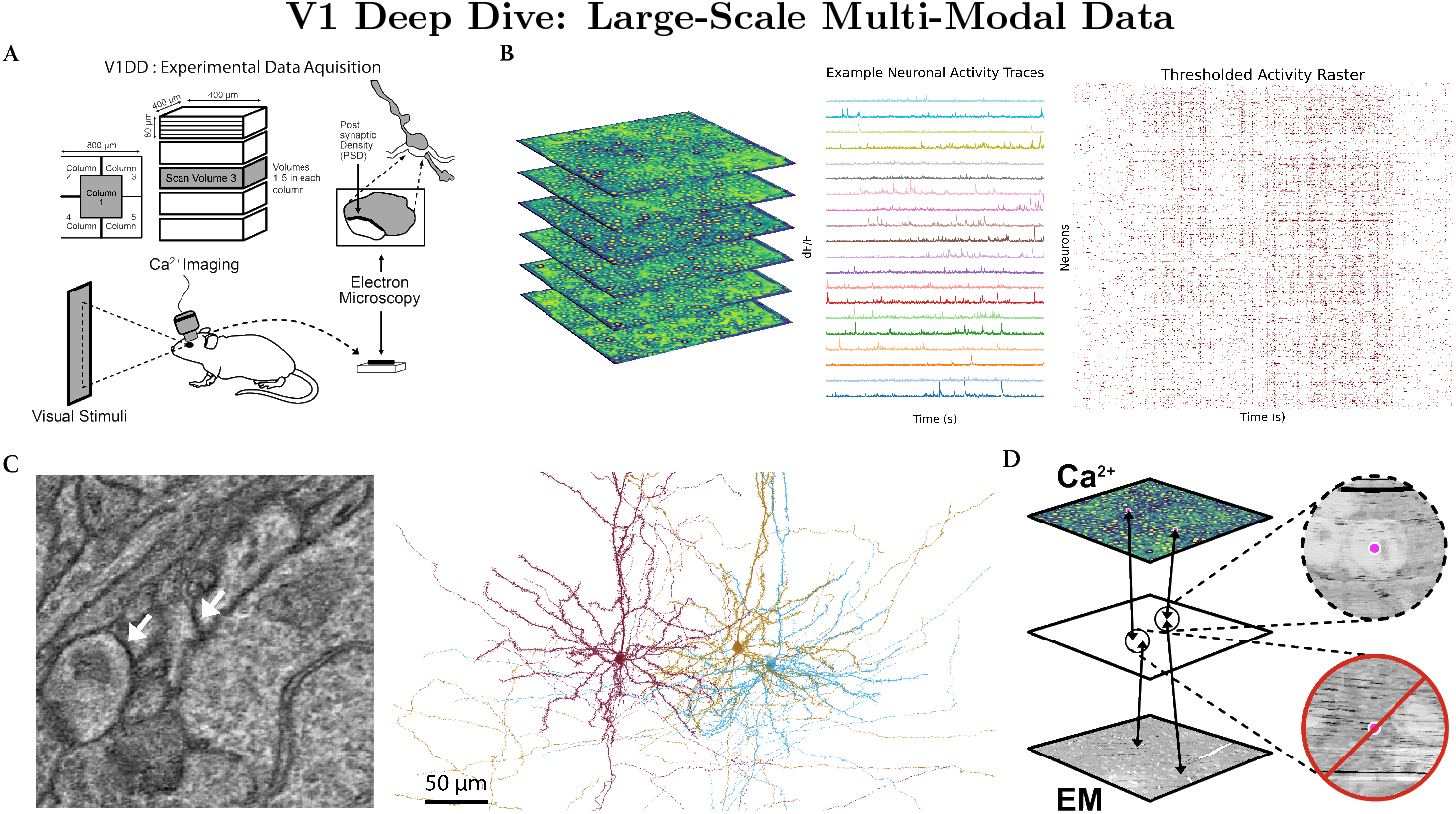
**(A)**. Schematic of experimental data acquisition for the V1 Deep Dive dataset (V1DD). V1DD consists of dense thousands of excitatory neurons in an 800 × 800 × 800 µm^3^ section of mouse V1, recorded during awake behaving imaging sessions, with no behavioral task. Our work focuses on the center column, particularly the third scan volume (pictured here in grey). Postmortem, the same tissue volume was fixed and imaged via transmission electron microscopy, allowing for reconstruction of synaptic connectivity, including post-synaptic density (PSD) volumes. **(B)** Each scan volume for in vivo imaging consisted of six stacked scan planes. Dense calcium activity allowed for the extraction of individual neuronal traces, with 20 example traces shown in addition to a raster plot of thresholded normalized activity for all 2708 neurons in Scan Volume 3 of Column 1. **(C)** Example of a microscopy view of connected neurons, with white arrows pointing to PSD. Reconstructed pyramidal cells corresponding to the left microscopy view are shown on the right. **(D)** Schematic showing the framework for coregistration of cells between the calcium recordings and the electron microscopy. Identified ROIs were mapped to an interstitial space (see Methods 5.1.4), where the correspondences were manually inspected.

These advances offer the opportunity to examine the structural correlates of Hebbian cell assemblies at an unprecedented scale. We extract assemblies from a Ca^2+^ fluorescence imaging scan and examine the reliability of their activation and their functional significance in the encoding of visual stimuli. We then analyze the connectivity between neurons based on cells that were coregistered between fluorescence and structural EM scans (Fig. 1D). Deriving hypotheses directly from postulates advanced by Hebb, we test his predictions about synaptic sizes and connectivity within and across assemblies, producing results that suggest a significant role for inhibition in their formation and activation, different from what is traditionally assumed.

## 3 Results

### 3.1 Neuronal Organization of Hebbian Assemblies

We analyzed a scan of the optical imaging dataset consisting of 2708 excitatory neurons recorded in parallel. The Similarity Graph Clustering (SGC) algorithm (Fig. 2A) generated 15 assemblies, which we ordered by size with ‘A 1’ representing the largest assembly (*n* = 1016) and ‘A 15’ the smallest (*n* = 23). A subset (*n* = 748, 27.6 percent of 2708 total) of neurons was assigned to no assemblies.

**Figure 2.**
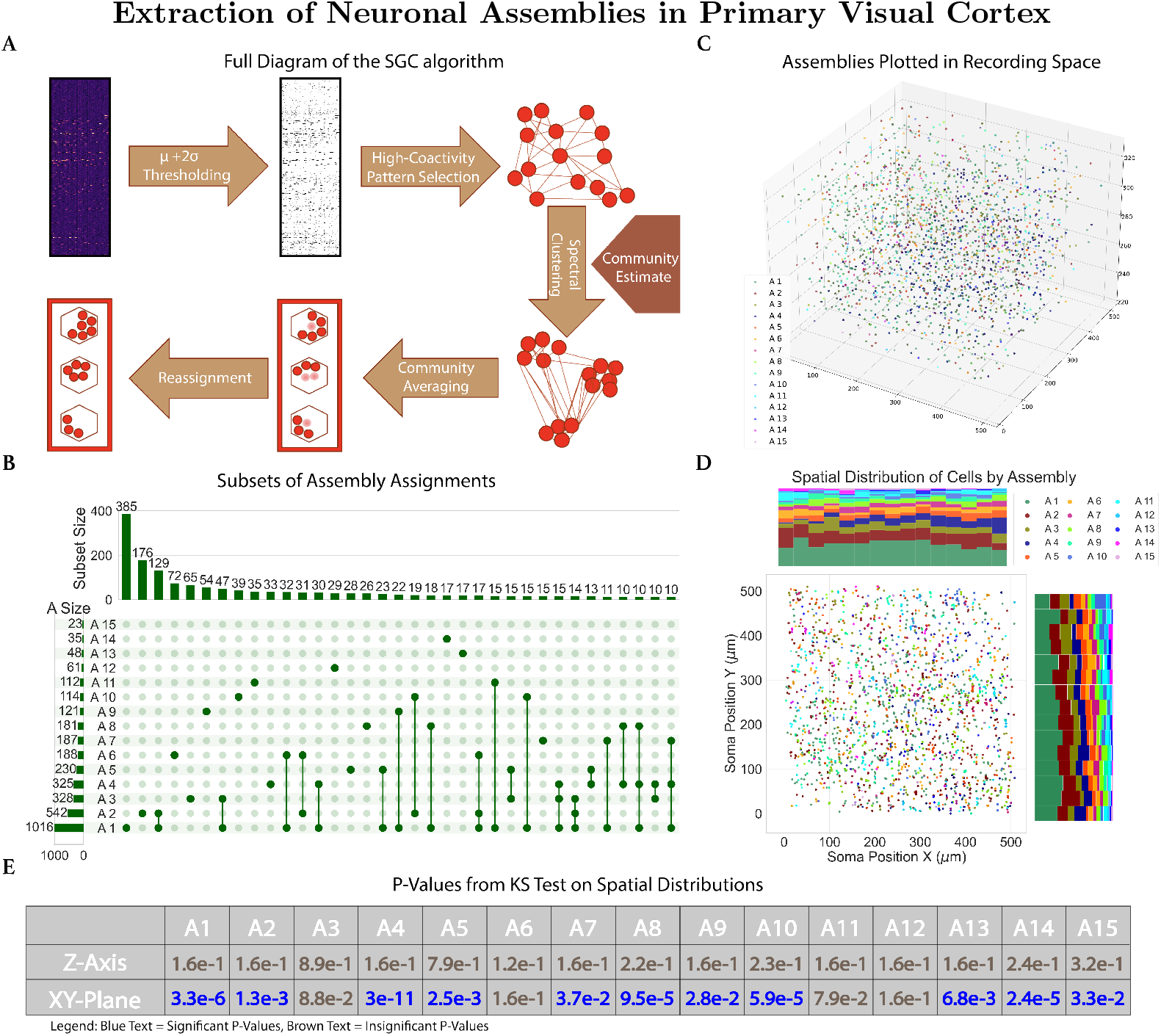
**(A)**. SGC, an extraction algorithm for assemblies uniquely designed for calcium imaging data, groups frames of the calcium fluorescence input to determine when neurons in assemblies are coactivated. Figure adapted from Mölter et al. [32] **(B)**. An UpSet [33] visualization of the subsets formed between assembly assignments. The histogram on the left represents the size of each individual assembly. The top histogram represents the size of the subsets between assemblies. Only subsets of ten neurons or greater were visualized. **(C)**. Spatial positions of fifteen extracted assemblies in the three-dimensional recording field. There are 1960 neurons visualized, including neurons assigned to multiple assemblies (plotted only once) but not including the 748 neurons that were assigned to no assemblies. **(D)**. Spatial distribution of assembly cells projected onto the x-y plane. Histograms for each axis are normalized to provide a per-bin proportional stack of the assembly distributions. **(E)**. A table presenting the KS test results on each assembly’s spatial distributions. Values colored in blue represent significant results (p-value *<* 0.05), while brown signifies insignificant results. All values have been corrected for false discovery under the Benjamini-Hochberg Procedure.

Their spatial distributions are shown in (Fig. 2C, D). Although spatial organization is not clearly visible, statistical analysis using the Kolmogorov-Smirnov (KS) [30] test revealed significant differences in the spatial organization of assemblies compared to that of the entire neural space (Fig. 2E). In particular, there was marked organization in most assemblies along the x-y plane, indicative of the retinotopic organization observed in studies of the visual cortex [31].

Most, but not all, assemblies shared members with other assemblies (Fig. 2B). For example, neurons assigned to ‘A 1’ were also assigned to ten other assemblies, highlighting Hebb’s proposal that precisely timed phase sequences allow a set of shared member neurons to participate in multiple assemblies.

### 3.2 Correlation and Sparsity

By construction, assemblies are expected to respond to distinct stimulus features, with a coactivity correlation sufficiently low to prevent their merging into a single assembly. Pearson correlation coefficients between assembly coactivity traces were significantly lower than the coefficients between coactivity traces of random ensembles of the same size distribution (see Methods) (Fig. 3A). As expected from traces derived from the average of population raster activity, both population groupings held significantly higher correlations than individual cell activity raster correlations, regardless of the subset of pyramidal cells being considered (assembly, non-assembly, or all individual cells).

**Figure 3.**
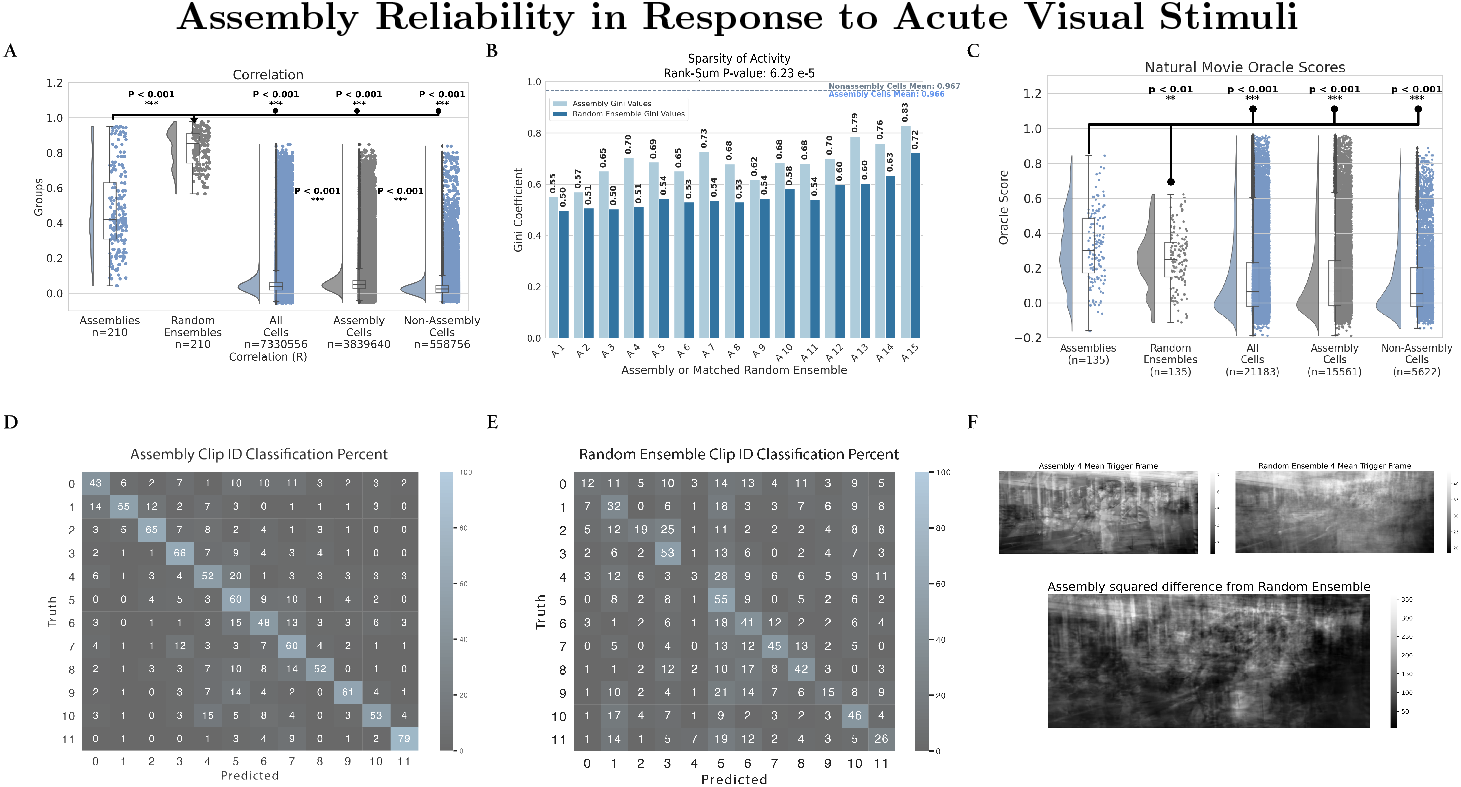
**(A)** Raincloud plot of pairwise correlations of coactivation between assemblies, size-matched random ensembles, and sets of individual cells. Coactivation for an individual cell is equivalent to a binary thresholded activity raster. **(B)**. Grouped bar plot of sparsity (measured by the Gini Coefficient) of coactivity over time in cell assemblies and the null-grouping of size-matched random ensembles. The random ensembles’ coefficients are significantly smaller than the set of assembly coefficients (Wilcoxon Rank-Sum p-value: 6.234*e* − 5). The average sparsity of individual assembly cells and non-assembly cells is also plotted as nearly equal horizontal dashed lines. **(C)**. Raincloud plot illustrating the reliability of activity from assemblies and general neuronal populations in response to natural movies. Oracle scores of each assembly and random ensemble coactivity trace were plotted, as well as the scores of sets of individual cells. These oracle scores are computed for the concatenation of natural movie clips and their responses, rather than individual clips, in order to reduce the likelihood of sparse responses causing an artificially high reliability score. **(D, E)**. Heatmap illustrating decoding accuracy of natural movie clips with Assemblies and Random Ensembles. Heatmap values indicate the accuracy of clip decoding by the percentage of presentation. Clip IDs, indicating a unique natural movie clip, are balanced such that each clip has an equal frequency of presentation. Values in the assembly heatmap are significantly greater than the random ensemble heatmap (Mann-Whitney u-stat: 7546.0, p-value: 6.07*e* − 5; one-sided on diagonal elements u-stat: 131.5, p-value: 3.25*e* − 4). **(F)**. Example plots of the mean ‘trigger frame’ of assemblies and random ensemble during natural movies. Frames were generated by averaging the frames associated with peak coactivity. The natural movie frame was visually better reconstructed by the assembly activity than that of the random ensemble, as signified by the plotted squared difference.

To further characterize the functional properties of assemblies, we computed the Gini coefficient [34] for each assembly’s activity trace. The Gini coefficient, a statistical measure of the ‘inequality’ of signal activity throughout the optical recording, revealed that assemblies exhibited highly sparse activity patterns (Fig. 3B), exhibiting significantly higher Gini coefficients than random ensembles (p-value : 6.23*e* − 5). The coefficient for each assembly ranged from 0.55 to 0.83, with particular assemblies with extreme sparsity, such as ‘A 13’ (0.79) and ‘A 15’ (0.83), exemplifying a high degree of functional selectivity. Interestingly, this sparsity metric was not solely dependent on assembly size, as intermediate-sized assemblies (e.g., ‘A 4’ through ‘A 12’) all exhibited similar Gini coefficients of around 0.70.

### 3.3 Assemblies Reliability in Stimuli Response

A section of the visual stimuli presented to the mouse consisted of natural movies. Across the hour scan time, twelve unique 15-second movie clips were repeatedly shown eight times. To evaluate the functional reliability of neuronal assemblies, we analyzed their responses to these stimuli. Reliability was quantified using an Oracle score, a leave-one-out correlation metric (see Methods 5.3) that measures the consistency of an activity trace across repeated presentations of the same stimulus. Oracle scores are used to measure the reliability of neuronal responses [35]. To provide a baseline with which to compare the functionality of these assemblies to tune to visual stimuli, the oracle scores of assemblies’ coactivity traces were compared to the oracle scores of all neuronal traces. In addition, we provide a population-level comparison with random ensembles.

Assemblies exhibited significantly higher Oracle scores compared to the average reliability of individual neurons (p-values *<* 0.0001), indicating that the assemblies, as populations, respond more consistently to visual stimuli (Fig. 3C). To ensure this result was not merely due to the inclusion of highly reliable neurons within assemblies, we separately calculated Oracle scores for neurons within assemblies and those assigned to no assemblies. We observed no significant difference between the cellular sets (p-values *>* 0.25). This result suggests that the reliability of these assemblies is derived from their collective activity rather than from the reliability of individual members.

Since population coactivity is expected to be more reliable in the general case than individual neurons, we compared our assemblies to random ensembles (defined as in Methods 5.2). Assembly coactivity traces demonstrated higher Oracle scores than coactivity traces computed in the same way as the assemblies for size-matched random ensembles. Complementary results are seen in the reliability of responses to orientation gratings (see Supp. Fig. 6). Population activity during the presentation of these gratings was used to characterize reliability as a function of orientation [36].

Further analysis revealed consistent patterns of high assembly coactivity during specific visual frames of these natural movies. We defined ‘trigger frames’ as moments when assembly activity exceeded a baseline threshold (see Methods 5.2) and found that these frames were highly consistent across repeated stimulus presentations. Example mean trigger frame pixel values are presented in (Fig. 3F), along with those of the corresponding size-matched random ensemble, and the squared difference. Visualizations of these frames suggest that assembly activity responds to complex features in the natural movies. These collective results provide evidence of assemblies’ ability to serve as functional populations with reliable and specific responses to visual stimuli.

### 3.4 Decoding Responses from Acute Visual Stimuli

We also assessed the ability of assemblies to decode visual stimuli by implementing a classification framework comparing assemblies to random ensembles (Fig. 3D, E). We employed a Multi-Layer Perceptron Classifier (MLPClassifier) to evaluate how well each grouping could decode the identities of the twelve natural movie clips. These classifiers have been shown to be effective in academic and clinical settings, with high levels of accuracy and flexibility in available hyperparameters [37, 38].

The results of our classifier revealed that assemblies significantly outperformed random ensembles in accuracy. Heatmaps of classification accuracy for natural movie clip identities demonstrated that assemblies provided more reliable decoding across repeated trials. A Mann-Whitney U-test confirmed this finding, with assembly accuracy significantly exceeding random ensembles for both overall performance (u-stat: 7546.0, p-value: 6.07*e* − 5) and diagonal elements (one-sided, u-stat: 131.5, p-value: 3.25*e* − 4), indicating enhanced stimulus-specific decoding.

### 3.5 Structural Organization of Assemblies

Coregistration between recorded activity and EM data provides us with a unique opportunity to explore the structural underpinnings of assemblies in the visual cortex. By mapping neural structure and connectivity at a micrometer resolution, we investigated the anatomical communication and organization of neurons with at least one shared assembly membership (shared assembly cells) compared to those with disjoint membership.

The strength of connections between the two groups was measured by performing a Wilcoxon Rank Sum test to compare the connection weights (defined as the sum of PSD volumes for all synapses between two cells). A Chi-squared test of Independence was performed to determine whether the pairwise frequency of connections differed between cells with shared assembly membership and those with disjoint assembly membership. To investigate higher-order structural patterns, we performed similar tests on sets of inbound and outbound disynaptic chains, as well as subdividing based on whether the intermediate cell in each chain was inhibitory or excitatory. 5.11.2

Our initial analysis of first-order connectivity surprisingly revealed no significant differences in the probability of direct monosynaptic connections between shared and disjoint sets or the strengths of those connections (Fig. 4D,E). This finding is incongruous with predictions that assemblies are defined by densely interconnected excitatory neurons [1] and suggested that the defining structural characteristics of assemblies might lie beyond simple pairwise connectivity metrics.

**Figure 4.**
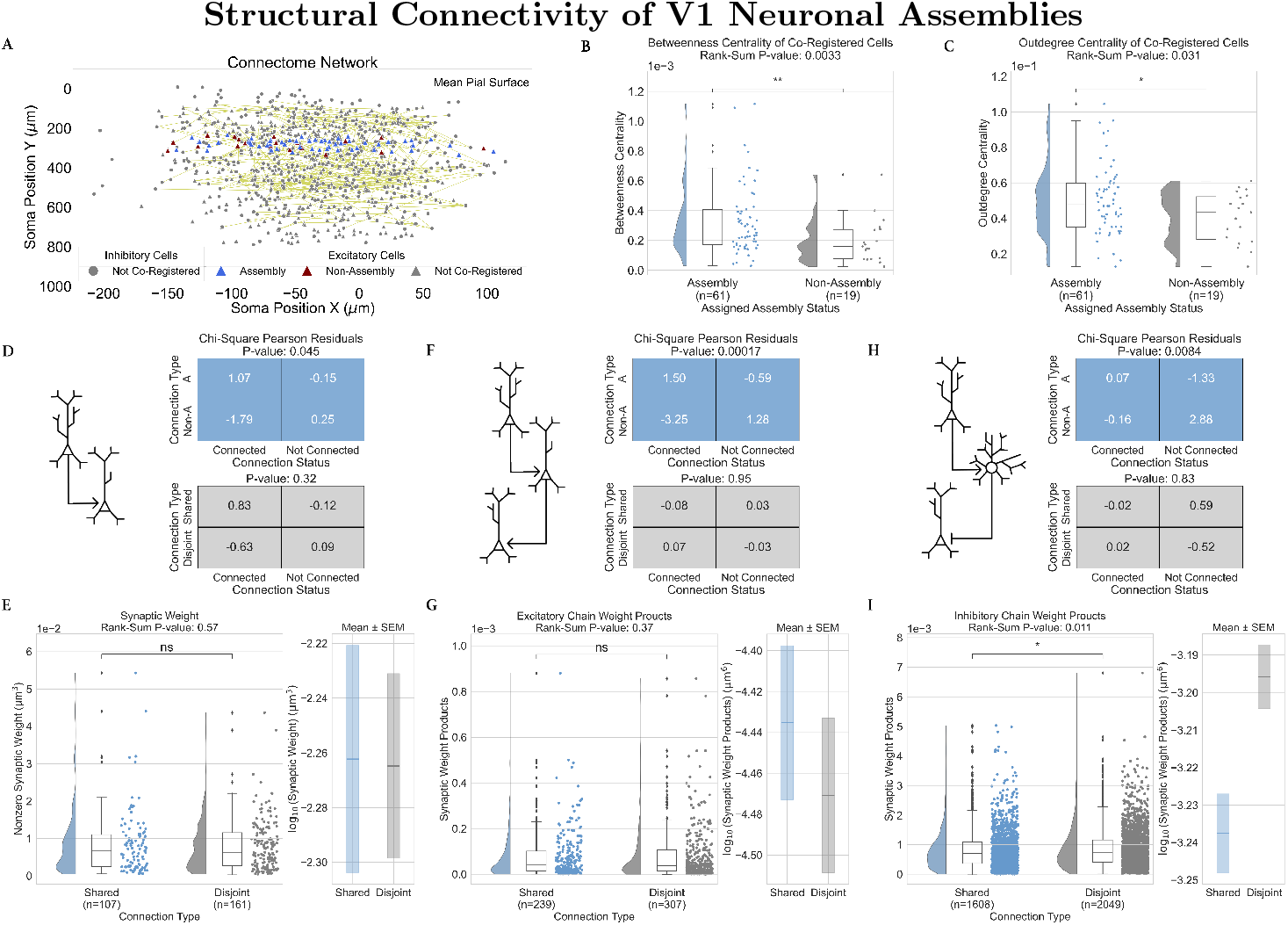
**(A)**. Visualization of the network being analyzed, showing the soma position of cells in the connectome, colored by cell-type and assembly assignment. All coregistered reconstructed neurons were found in layer 2/3 or layer 4 of V1. **(B)** A raincloud plot of betweenness centrality, demonstrating a higher centrality for assembly neurons than those not in assemblies(Wilcoxon Rank-Sum: p-value *<* 0.01). **(C)** A raincloud plot of outdegree centrality, a mathematical proxy for probability of connection, demonstrating a higher centrality for assembly neurons (Wilcoxon Rank-Sum: p-value *<* 0.05). **(D**,**F**,**H)**. Chi-squared analysis of the likelihood of monosynaptic (D), disynaptic excitatory (F), and disynaptic inhibitory (H) connections (schematic on the left). This comparison was made between assembly neurons and non-assembly neurons (top-right) as well as between neurons that share an assembly membership and neurons that participate in disjoint assemblies (bottom-right). **(E, G, I)** Raincloud plots showing the combined synaptic PSD volume per extant monosynaptic (E), disynaptic excitatory (G), and disynaptic inhibitory (I) connection, each divided between origin and terminus cell pairs which share assemblies and those which participate in disjoint assemblies. Log-scaled plotting of the chain weights with SEM-based confidence intervals is included as an inset plot to the right of each panel.

To investigate higher-order structural patterns of these assemblies, we conducted a motif analysis. In a neural network, when a significant number of subgraphs containing a small number of interconnected cells repeat a particular pattern (e.g., Cell Type A → Cell Type B →− Cell Type A), they are classified as a motif [39, 40], with each subgraph counted as a motif instance. The frequency with which these motifs persist has been predictive of correlation in similar neural networks [41].

Our analysis concentrated on second-order chain motifs, or structures consisting of three neurons connected by two synaptic links (schematic shown in Fig. 4F,H). Second-order neural motifs are divisible into various types based on the arrangement of their synaptic connections. For this study, we prioritized chain motifs, as they allowed us to utilize aspects of the EM dataset that are not yet coregistered to activity recordings for the central elements in the chain, so long as the first and last cells in the chain are coregistered.

Notably, while our excitatory chain analysis revealed no significant differences between shared and disjoint assembly memberships (Fig. 4F,G), an analysis of inhibitory motifs convey a different story: disjoint assembly memberships exhibited significantly stronger feed-forward inhibitory connections than shared memberships (Fig. 4H,I). Log-scaled plotting of the chain weights (Fig. 4I, inset) revealed a complete lack of overlap between SEM-based confidence intervals, suggesting that this is a result with both reliability and a non-trivial effect size. Notably, this result was insignificant when restricting our analysis to only the inhibitory synapse or only the excitatory synapse within the feed-forward inhibitory chain, suggesting that both connections may play a role.

The significantly stronger feed-forward inhibitory connections suggest a mechanism of mutual inhibition that may regulate the interaction between distinct assemblies, preventing excessive coactivation and ensuring the discriminability of assembly responses to inbound information. Indeed, we find a negative Pearson r correlation coefficient between the disjoint feed-forward inhibitory chain weights and the correlation of the two disjoint assemblies’ coactivity traces (r statistic : −0.19, p-value : 0.0048; see Methods 5.11.4).

All together, our structural results reveal that assemblies are not anatomically defined solely by a fundamental increase in local connectivity, but through their higher-order patterns of organization. Stronger feed-forward inhibitory connections in disjoint memberships suggest a possible mechanism for maintaining functional segregation between assemblies. These structural insights provide intuition on the mechanistic basis that drives the functional properties of assemblies, reinforcing their contribution as modular units of sensory processing.

### 3.6 Non-Assembly Cells

The same monosynaptic and disynaptic analysis run on pairs of cells with shared assembly memberships and cells with disjoint memberships was also run on pairs of assembly cells (cells that both participate in at least one assembly) and pairs of non-assembly cells (cells that had no assembly memberships for either cell).

We show assembly cells exhibiting a pattern of connectivity distinct from non-assembly cells, a requirement of Hebb’s theory. We assessed the higher-order integration of assembly neurons into the broader structural network using centrality metrics. In particular, we found significantly lower betweenness centrality (Fig. 4B), a measure of a node’s importance in mediating communication within a network, and outdegree centrality (Fig. 4C), an analog for probability of outbound connectivity, in neurons outside assemblies compared to neurons within assemblies. Furthermore, we found that the pairwise probability of monosynaptic connections, disynaptic excitatory chains, and disynaptic inhibitory chains was significantly greater for cells sharing assembly membership than for non-assembly cells (p-values : 0.045, 1.7*e* − 4, and 0.0084, respectively). No significant differences in PSD Volumes between assembly and non-assembly cells were observed. Importantly, these results were not due to differences in the spatial locations of assembly and non-assembly cells with respect to the centroid of the network (Supp Fig. 9). These findings confirm that cells in the connectome participating in at least one assembly are more interconnected than those not participating in any.

Overall, these results indicate reduced participation in the structural framework of the primary visual cortex for non-assembly cells and imply an organizational role of assembly neurons as hubs for information flow. In contrast, non-assembly neurons may play a more peripheral, secondary role in the network, such as noise filtering.

## 4 Discussion

### 4.1 Inhibition as the Delineating Basis of Hebbian Assemblies

This work provides the first demonstration of the structural underpinnings of Hebbian assemblies and validates Hebb’s assembly theory in a surprising way; while there is strong agreement on functional predictions, our structural findings run counter to Hebb’s original proposal. We do not observe a marked increase in the strength or probability of excitatory connections between cells that participate in the same assembly (Fig. 4D-G). Instead, we find significantly stronger feed-forward inhibitory connections targeting cells in other assemblies (Fig. 4I). Hebb and others [42, 9, 3] have hypothesized that targeted inhibition could play a crucial role in the formation and delineation of assemblies. Our findings offer the first clear empirical evidence for this mechanism.

Despite this novel result, our findings remain a validation of Hebb’s structural postulates and are consistent with what would be required for assemblies to be delineated by inhibition. Assembly cells were substantially more integrated into the connectome, displaying significantly higher betweenness and outdegree centrality (Fig. 4B,C). This pattern demonstrates that cells participating in our activity-defined assemblies possess markedly different patterns of connectivity from cells that do not, establishing the necessary structure-function relationship postulated by Hebb.

Intuitively, the delineation of assemblies can be realized on a spectrum between internal (shared-assembly) excitation and external (disjoint-assembly) inhibition. Shared-assembly excitation would rely on broad inhibitory connectivity across the entire population of assembly neurons, and disjoint-assembly inhibition would similarly rely on broad excitatory connectivity. We find a significantly greater probability of excitatory connections between cells belonging to any assembly (Fig. 4D,F), however we did not observe significantly stronger or more frequent excitatory connections between neurons with shared assembly membership (Fig. 4D-G). Combined with our finding of significantly greater feed-forward inhibitory weights between neurons with disjoint assembly membership (Fig. 4I), the broad excitation strongly suggests that the extracted assemblies are delineated predominantly via inhibition.

### 4.2 Functional Consistency of Assemblies

Functionally, ensembles of cells classified as assembly cells behave in ways consistent with Hebbian assemblies. The assemblies demonstrated markedly higher reliability in their responses to naturalistic visual stimuli compared to individual cellular responses of all three sub-populations of cells and size-matched random ensembles, reflected by significantly elevated Oracle scores (Fig. 3C). This consistent coactivity of the assemblies has been hypothesized to offer resilience to representational drift, allowing them to serve as a substrate for long-lasting representation [8]. Representational drift reflects a functional characterization of fluctuations at the cellular level, such as synaptic turnover [43, 44], even under stable stimulus conditions [45]. This phenomenon has been observed in areas other than V1, such as the piriform cortex [46]. By maintaining consistent and coherent patterns of coactivity, assemblies may offer a general cortical mechanism of stable perceptual representations despite such turnover.

Beyond their reliability in responses, assemblies were also superior in our decoding of natural movie presentations (Fig. 3D), underscoring their efficacy in extracting higher-order visual information. In addition, the sparsity of assembly coactivity was substantially greater than that of random ensembles (Fig. 3B), which, when combined with a lower average correlation 3A), is consistent with assemblies employing a cost-effective encoding strategy [47]. Such a strategy would enhance the capacity for distinct representation of sensory inputs, translating more flexible individual responses into reliable population-level encoding. Finally, cells assigned to assemblies exhibit significantly higher pairwise correlations than non-assembly cells, corroborating the experimental work of Harris and Carandini [48] defining ‘choristers’ and ‘soloists’. However, we found no difference in their average response reliability to natural movies or in their overall signal sparsity. This result implies that a population perspective is required for the encoding of reliable perceptually relevant stimulus features.

### 4.3 Bridging Structure and Activity

Some examined ensemble features bridge structure and activity, a crucial aspect of this study, which has not previously been achievable at scale. First, the extensive overlap of neurons across multiple assemblies (Fig. 2D) is consistent with both Hebb’s proposal of sub-assemblies [9] and Yuste and colleagues’ review of definitions put forward for ensembles [49], with individual neurons contributing to multiple functional modules. Second, our observation of greater chain weights in feed-forward inhibitory chains offers a mechanistic explanation of how assemblies can retain distinct responses to stimuli without becoming so correlated in their activity that they merge into a single functional ensemble. The plausibility of this explanation is supported by the significant observed negative correlation between inhibitory chain connection strength between disjoint assemblies and the r coefficient of their paired assembly coactivity traces, showing that the greater the inhibitory weight of the connections observed, the less correlated the activation of the two assemblies is 3.5.

### 4.4 Further Directions

We observe that the assemblies exhibit notable variability in size (Fig. 2D). SGC minimizes overestimation of the number of assemblies or neuron assignments, suggesting that these size differences reflect intrinsic properties rather than methodological artifacts. There is therefore an opportunity for future studies to delineate the distinct roles potentially served by larger assemblies, such as ‘A 1’, compared to their smaller counterparts. In addition, while novel in its scale and multifaceted nature, we see opportunities to reduce some of the current limitations within this dataset. Continued reconstruction, proofreading, and coregistration will allow analyses of more neurons and subsequent new lines of inquiry. For instance, the SGC algorithm additionally flags frequent patterns of activity, potentially analogous to Hebbian phase sequences [6, 7]. We currently do not have sufficiently many coregistered and fully reconstructed cells to allow examination of connections that bridge one pattern to the next. Exploring how these patterns interact could reveal mechanisms of large-scale neural coordination. Other limitations will be more difficult to overcome without the availability of next-generation multi-modal datasets. Most importantly, the temporal resolution of the scans leaves us with no ability to examine activity on a tens-of-ms timescale, which is highly relevant for synaptic plasticity.

These results suggest several potential implications for the field at large. Hebb proposed assemblies as a universal building block, simultaneously addressing the functional and structural sides of perception, cognition, and behavior. A basic unit that bridges structure and function allows one to derive structural predictions from functional characterizations and vice versa. Beyond validating Hebb’s theory, the evidence presented here for an underlying inhibitory mechanism provides further support for the analysis of inhibition in cognitive and sensory disorders studied to date primarily through the lens of excitation [50, 51]. In the future, the integration of cell-type-specific genetic information with functional assembly data could also provide deeper insights into the molecular foundations of assembly formation and maintenance in health and disease [52]. Hebb postulated that assemblies form the atoms of cognition, and it has not escaped our notice that the cross-inhibitory mechanism we here demonstrate might be a universal feature of brain-wide assembly organization.

## 5 Methods

### 5.1 V1 Deep Dive Dataset

#### 5.1.1 Stimuli

Visual stimuli were presented using the same monitor configuration as in de Vries et al. [53]. Imaging sessions were one hour long and offered a wide variety of visual stimuli. Assembly extraction was performed on the fluorescence data from the full session. The remainder of our analysis utilizes only the natural movie clips and the full-field drifting gratings, details of which are provided below. The other stimuli presented during the session included natural images, windowed drifting gratings, and locally sparse noise. The natural movies stimulus consisted of a series of clips concatenated into 3,600 frames (with 30 Hz frame rate), presented 8 times.

The full-field drifting gratings stimulus consisted of a drifting sinusoidal grating at a 1 Hz temporal frequency and 80 percent contrast, presented at 12 different orientations (multiples of 30°) and at 2 spatial frequencies (0.04 and 0.08 cycles per degree). Each condition was presented eight times, in randomized order, with one second of mean luminance grey between presentations. The windowed drifting grating stimulus matched the full-field stimulus, but the stimulus was restricted to a 30° diameter window. For each column, the position of the window was determined separately to align with the population receptive fields of imaged neurons. The locally sparse noise stimulus consisted of white and dark spots on a mean luminance gray background. Each spot was a square 9° on a side. Each frame of this stimulus was presented at 3 Hz.

Two natural scene stimuli were presented. One consisted of 12 images (selected from those used in the Brain Observatory pipeline) presented in a frozen sequence, and repeated 40 times. The other consisted of the full 118 images from the Brain Observatory pipeline, presented 8 times total. The images were presented in a random order, but fixed with two different seeds. Each of the two seeded sequences was presented 4 times. The images were presented at 3 Hz.

#### 5.1.2 2/3 Photon Microscopy and Activity Data Processing

The full 800 × 800 × 800 µm^3^ volume was divided into five columns, each imaged via either 2-photon (2P) or 3-photon (3P) microscopy, depending on depth. The volumes were scanned over the course of several sessions. Each session consisted of the full set of visual stimuli (see below), presented in the same order and with the same timing. Our analysis is concerned only with the central column, centered within the 800 × 800 × 800 µm^3^ volume, where the largest set of reconstructed, coregistered, and proofread neurons is currently available. Figure 1A shows the arrangement of scan volumes (5 volumes spanning 75 µm to 620 µm depth, each 400 × 400 × 80 µm^3^) and imaging planes (6 planes within each scan volume, separated by 16 µm at thirty-seven frames per second so that each plane is imaged at 6 Hz.) We selected volume three of the central column for our primary analyses and validated our work with volume four. Both were imaged with 2P microscopy.

The fluorescence data was preprocessed using the standard LIMS pipeline as used for the Allen Institute’s Visual Coding 2P dataset [53], including motion correction, segmentation, demixing, neuropil subtraction, ROI filtering, and df/f calculation. Identified regions of interest were run through a classifier trained on manual labeling data meant to reduce the false classification of artifact ROIs as neuronal somas, and only those with a high confidence score (at least 0.5) were included in our analysis.

#### 5.1.3 Electron Microscopy and Reconstruction

The mouse was transcardially perfused with a fixative mixture of paraformaldehyde and glutaraldehyde. All procedures were carried out in accordance with the Institutional Animal Care and Use Committee at the Allen Institute for Brain Science. The largevolume staining protocol was adapted from [54]. After dissection, the neurophysiological recording site was identified by mapping the brain surface vasculature. A thick (1200 µm) slice was cut with a vibratome and post-fixed in perfusate solution for 12 to 48 h. The tissue was then infused with heavy metals, dehydrated, and embedded in EMS Hard Plus resin. After curing, the samples were epoxy cured to a stub. They were then sliced and placed onto continuous tape by a ATUMtome Automated Tape Collecting Ultramicrotome.

The continuous tape was fed into an automated high-throughput transmission electron microscopy pipeline[55]. Transmission electron microscopy is particularly well suited for automated imaging and preserves very good x-y resolution at the expense of some resolution on the vertical z-axis, and so specialized methodology was deployed during reconstruction[56, 57]. Serial section alignment was performed through a contract with Zetta A.I, followed by stitching [58, 55], segmentation, and automated reconstruction. Proofreading of a subset of cells was performed under contract by Ariadne.ai.

Cell-type predictions were made for single-nucleus objects within 175 microns of the centerline and with a nucleus volume greater than 218, based on dendritic skeleton features adapted from [59]. Segmentation and annotation were stored in a CAVE database for access via CAVEClient [60].

#### 5.1.4 Coregistration

Manual coregistration was performed using the Fiji plugin BigWarp [61, 62]. A structural scan of the vasculature was aligned with the two-photon imaging planes (max intensity projection). Next, a downsampled EM image was aligned to this composite using a thin-plate-spline transform based on manual landmarks. After initial alignment, the transform was used to predict additional correspondences between two-photon ROI centroids and segmented EM cells. Four hundred verified correspondences between a fluorescence imaging ROI with a high classifier score and a corresponding morphologically typed EM-reconstructed cell passed manual inspection by two independent reviewers and were included in this study. 315 of these correspond to cells whose fluorescence was recorded in the scan we analyzed (volume three of the central column). Of these 315, 80 had axon reconstructions that were verified accurate by trained experts (‘proofread’) to their maximal extension within the scan volume. Coregistration and reconstruction are ongoing at the time of this writing, but the extant data already allow for a relation of the physiology of neural data to its exact anatomy at an unprecedented level.

### 5.2 Graph Clustering for Assembly Extraction

To extract assemblies, we use the Similarity-Graph-Clustering (SGC) algorithm that has been originally proposed for the detection of neural assemblies during spontaneous activity in the zebrafish optic tectum [63]. The SGC algorithm identifies neuronal assemblies using ideas from graph theory by transforming the problem into one of community detection on some graph in which assemblies correspond to distinct (graph) communities. Unlike traditional methods that rely on pairwise correlations between cells, SGC groups frames of fluorescence indicative of significant coactivity [32]. These moments of higher-order correlation are referred to as potential ‘phase sequences’, representing when populations of neurons act cohesively as a closed circuit during assembly activation [64].

Importantly, by design, SGC allows cells to participate in multiple assemblies. Assembly overlap has been integral to assembly studies in the past [65]. Hebb originally postulated that the shift between active assembly states could be what is colloquially referred to as a ‘train of thought’ [1]. Our results revealed a sizable degree of overlap between assemblies, which was visualized through an UpSet plot (Fig. 2D) [33], showing neurons frequently assigned to multiple assemblies (Fig. 2D).

Compared with several other prominent assembly extraction algorithms upon application to both simulated and biological calcium imaging datasets under different conditions, this algorithm was shown to perform best overall [32]. Although it did not yield a perfect reconstruction of the assemblies in the biological dataset, SGC was able to recover the assemblies with higher accuracy than all other algorithms. In part, this performance has been attributed to the computational effort that SGC places in estimating the number of assemblies before defining them.

We used a recent implementation of the SGC algorithm in Python [66]. For completeness, we briefly recall the main steps here: The algorithm begins by thresholding the calcium fluorescence signals (df/f) of the ROIs (neurons) by two standard deviations above the mean to minimize noise (Fig. 2B). Afterward, activity patterns are selected where the coactivity level of neurons exceeds the significance threshold. This threshold that determines the set of high-activity patterns is based on a null model of coactivity obtained by shuffled activity signals (significance value: 0.05, rounds: 1000). A *k*-nearest-neighbor graph is then constructed from the set of high-coactivity patterns based on the similarity between the patterns in the cosine distance. For that, the number of neighbors *k* is automatically chosen such that the resulting graph is connected. In the next step, the number of communities in this graph is estimated using a statistical inference procedure. For our study, we fixed the hyperparameters with five independent Monte Carlo rounds of 150, 000 steps each. With an estimate for the most likely number of communities, spectral clustering is applied. These clusters of high-coactivity patterns are the first prospective selection of coactivity patterns corresponding to the assemblies. However, the final step is a combination of averaging and reassignment to reject groups that may have been erroneously defined because of a high level of noise in the original signal. This minimizes the likelihood of overestimating the number of assemblies or the neurons that should be assigned to those assemblies. Neurons are assigned to assemblies based on their affinity, the probability that they were active in any of the assemblies’ activity patterns (affinity: 0.4). To establish our hyperparameters, we first performed a grid search on an independent functional scan volume (Scan Volume 4, acquired under identical conditions). Candidate values of Monte Carlo rounds for our estimation step (50000, 100000, 150000) and affinity cut-offs (0.2, …0.3, 0.9) were evaluated by how well the resulting assembly assignments could fit to a low-dimensional embedding of pairwise activity correlations (see Supp. Fig. 7). TO compare the regression fits, we applied the Akaike Information Criterion, which is calculated from the number of independent variables and the maximum likelihood estimate of the model. The parameters that maximized the Akaike Information Criterion in the regression fits of scan volume 4 were 150000 Monte Carlo rounds and an affinity cut-off of 0.4. These parameter values were then locked and applied to scan volume 3.

### 5.3 Random Ensembles

Random ensembles served as a null model and were defined as randomly selected sets of neurons drawn with equal probability and no replacement from the population of all recorded pyramidal cells within scan volume 3. Each set was size-matched to its corresponding assembly, with the same number of neurons.

### 5.4 Oracle Scores

Oracle scores are a measure of the reliability of the trace response of cell activity to repeated visual stimuli computed through a jackknife mean, or leave-one-out mean [67, 68], of correlations between the activity trace across the repeated visual stimuli. In this work, we treat the entire stereotyped stimulus sequence as a single stimulus, concatenating natural movie clips and their responses, rather than calculating the score for individual clips. This should reduce the risk of sparse responses leading to an artificially high reliability score.

Let *S*_*i*_(*t*) denote the activity trace of a neuron or an assembly during the *i*-th presentation of a stimulus at time *t*. Let 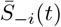 represent the mean activity trace of all activity traces excluding the *i*-th presentation, out of *n* total presentations of the same stimulus. From this, we can calculate the Oracle score *O* as

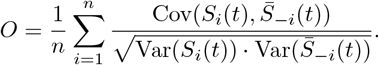

In this formulation, the numerator will calculate the covariance between the activity trace at each presentation to the mean activity of all other repeats, while the denominator scales the magnitude of this covariance by the product of the standard deviations of *S*_*i*_(*t*) and 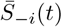.

### 5.5 Trigger Frames

To assess the visual stimuli associated with high activity in neuronal assemblies, we computed trigger frames by identifying peak activity times and extracting the corresponding images from a natural movie presentation. From a coactivity trace for assembly *k* over time *S*_*k*_(*t*), we can detect peaks *P*_*k*_ using the `scipy` signal package to define local maxima. We then define a mean trigger frame *µ*_*k*_(*x, y*) for a pixel (*x, y*) as

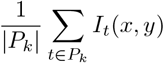

where *I*_*t*_(*x, y*) is the natural movie frame at time *t*.

This allowed us to visualize the average triggering frame for each assembly. The same process was then repeated for the coactivity traces of the size-matched random ensembles, and the squared difference between the average frame for each assembly and the average frame for its corresponding random ensemble. These computations allowed us to determine which frame of visual stimuli and consistent features in those stimuli were most strongly associated with high assembly coactivity in particular, not merely broad neuronal activation.

### 5.6 Decoder

To evaluate the ability of assemblies to decode visual stimuli, we implemented a Multi-Layer Perceptron Classifier (MLPClassifier) from the `scikit-learn` library. This classifier was used to differentiate between 15 natural movie clips based on assembly coactivity time traces. Random ensembles of neurons with the same size distribution as the assemblies were used as a null model for comparison.

The MLP is a classical feed-forward neural network composed of an input layer, one or more hidden layers, and an output layer. Each node in a layer is fully connected to every node in the subsequent layer through weighted connections. The final output of the function is determined by a non-linear activation function applied to the weighted sum of its inputs plus a bias term.

The MLPClassifier from `scikit-learn` is an implementation of a Multi-Layer Perceptron (MLP), a type of feed-forward neural network. It operates by mapping input features **x** to outputs **ŷ** through a series of hidden layers. Each layer consists of neurons that perform a weighted sum of their inputs, followed by the application of a non-linear activation function. Defining **W**^(*k*)^ as the weight matrix for the *k*-th layer, **h**^(*k*)^ as the input, and **b**^(*k*)^ as the bias vector for the *k*-th layer, we can define with activation function *σ*

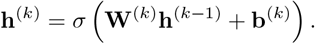

For classification, the final layer uses the softmax activation function to output probabilities for each class. With *L* denoting the number of layers, we can define these output probabilities as

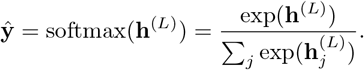

The MLPClassifier is trained using backpropagation, optimizing the weights and biases via stochastic gradient descent or adaptive solvers such as Adam. Regularization can be applied through an 𝓁_2_-penalty term, controlled by a hyperparameter.

The assembly coactivity time traces were paired with corresponding natural movie clip IDs. The data was split into training and test sets using an 80-20 split, and features were scaled to normalize the input. A cross-validated grid search was used to optimize the hyperparameters.

### 5.7 Gini coefficient

The Gini coefficient [34], a statistical measure that exemplifies the state of inequality within a population. While often applied in economics to evaluate income inequality, this metric has been applied as a valid approximation for signal sparsity [69, 70]. This rendition of the application has been shown to serve as a relatively simple and robust measure [71]. For our study, the coefficient is employed to quantify assembly signal heterogeneity.

The Gini coefficient, *G* is often calculated with respect to the Lorenz curve, which plots the cumulative distribution of a set (e.g., assembly coactivity trace) against its rank in ascending order. For a given assembly *A* with coactivity trace *S* = [*s*_1_, *s*_2_, …, *s*_*T*_] where *s*_*t*_ is the proportion of active neurons at time point *t*, the coefficient for that assembly is then calculated as

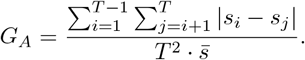

This computation is performed independently for each assembly, providing a metric of signal inequality. A value of 0 for *G*_*A*_ implies all values are identical, while a value of 1 indicates perfect inequality *i*.*e. in a time series a single time point contains all the activity*. A high value of *G*_*A*_ indicates that coactivity is dominated by a small number of time points, reflecting the temporal sparsity.

### 5.8 Correlation

We examine coactivity correlations for each pair of assemblies, each pair of random ensembles, and each pair of neurons within each of our three sets of neurons (see Fig. 3A-C). To do so, we computed the Pearson’s correlation coefficient r between the two coactivity traces (where the coactivity trace of a single neuron is mathematically equivalent to its thresholded raster activity, per [32]).

### 5.9 Motif Extraction

Motifs were extracted with the DotMotif Python package which detects subgraphs within a graph based on the principle of subgraph monomorphism. For every graph *H* = (*N*_1_, *E*_1_) given by the user, the Dotmotif algorithm detects subgraphs *G*^*′*^ within the graph *G* = (*N*_2_, *E*_2_) such that there exists a mapping 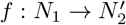 where 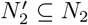 and for every edge (*a, b*) ∈ *E*_1_, the corresponding edge 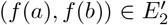. The matched subgraphs may also contain additional edges within the graph *G*. This algorithm was used to detect disynaptic chains within the connectome.

To further refine our analysis, chain motifs were classified according to the type of intermediary neuron, distinguishing between excitatory and inhibitory connections, with the latter providing insight into feed-forward inhibition mechanisms.

### 5.10 Motivating Postulates

Our statistical analyses involve tests of the following postulates. First, excitatory connections between cells that share at least one assembly (‘shared’ connections) will be stronger than connections between cells which do not participate in any of the same assemblies (‘disjoint’ connections), due to Hebbian plasticity [1]. This was examined both with regard to the post-synaptic density volume of monosynaptic connections between known coregistered cells within the dataset, and separately in the form of the product of connection PSD volumes in disynaptic excitatory chains which originated and terminated with shared or disjoint cells, allowing us to evaluate indirect excitatory connections where the middle cell had not yet been coregistered.

Second, that excitatory connections will be more frequent within assemblies than between assemblies, due to a combination of Hebbian plasticity and pruning of synapses. A number of computational studies have shown that long-term potentiation [72] and pruning [73] play important roles in effective Hebbian assembly formation. This was examined in the form of per-connection targeting statistics, as well as per-cell inbound and outbound probability of both monosynaptic and disynaptic shared and disjoint connections.

Third, as discussed in the introduction, Hebb himself suggested that the assemblies could have been formulated via modulation of inhibition. Additionally, a number of computational models [74] have relied on inhibition between assemblies to restrict simultaneous activation, enabling competition between assemblies. It was, therefore, taken as an established hypothesis that one would expect inhibition between assemblies to be greater than within a given assembly. As monosynaptic connections between excitatory cells cannot be used to evaluate inhibition, this postulate was examined only in disynaptic chains, where the middle cell was morphologically classified as an inhibitory interneuron. Our examination involved the product of connection PSD volumes in such disynaptic inhibitory chains bridging shared and disjoint assembly neurons, along with a per-cell evaluation of the inbound and outbound probabilities of disynaptic inhibitory chain connections.

Fourth, and finally, we acknowledge that Hebb discusses the reinforcement of sparse connectivity in his accounts of the emergence of cell assemblies [42], and thus this reinforcement might have a significant effect on a sparse subset of connections while producing a minimal difference in the central tendency of the overall set of connections. Dorkenwald et al. [75] demonstrated a bimodality of the log PSD volume of excitatory-excitatory synapses in the similar MICrONS mm^3^-dataset [26] (see also Supp. Fig.8 for result in V1DD), suggesting that a subset of such connections is impacted differently by processes determining PSD volume. Combining these two, we decided to test the hypothesis that the larger of the two-component distributions found in [75] would be more likely than chance to involve connections between shared assembly neurons.

### 5.11 Statistical Methods

In this section, we detail the motivations and specifics of our analysis and methods. We analyzed differences in connectivity metrics between shared-assembly and disjoint-assembly connection types, focusing on both the probability and strength of connections. Our analysis considered both direct monosynaptic and disynaptic connections between neurons (“by connection”), as well as the sets of inbound and outbound connections grouped by cell (“by cell”). For disynaptic chains, we grouped by whether the intermediate (middle) cell was excitatory or inhibitory.

To evaluate connectivity metrics, we first defined the connection types based on assembly membership. Let *A* be the set of all assemblies, with *A*_*i*_ denoting the subset of assemblies that include cell *i*. Formally,

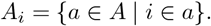

Using these subsets, we defined the following binary indicators to capture the assembly relationship between pre-cell *j* and post-cell *i*:

- Shared_*ij*_ = 1 if *A*_*j*_ ∩ *A*_*i*_≠ ∅
- Disjoint_*ij*_ = 1 if *A*_*j*_ ∩ *A*_*i*_ = ∅
- Assembly_*ij*_ = 1 if *A*_*j*_≠ ∅ and *A*_*i*_≠ ∅
- Non-Assembly_*ij*_ = 1 if *A*_*j*_ = ∅ and *A*_*i*_ = ∅

#### 5.11.1 Monosynaptic Metrics

We defined *w*_*ij*_ as the pairwise summed post-synaptic density (PSD) between pre-cell *j* and post-cell *i*, and *b*_*ij*_ as an indicator variable that takes the value 1 if at least one synapse exists between *j* and *i* and 0 otherwise. All metrics exclude autapases, as these were not reliably represented in the dataset (*j* ≠ *i*).

The probability of monosynaptic outbound connection for a pre-cell *j* was calculated as the proportion of realized connections under a given connection type C ∈ {Shared, Disjoint, Assembly, Non-Assembly}, normalized by the total number of potential post-cell partners for that connection type,

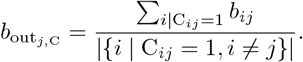

Similarly, the probability of a monosynaptic inbound connection for a post-cell *i* was defined as

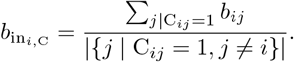

For connection strength, we computed the average realized summed monosynaptic outbound PSD for a pre-cell *j* as the total PSD across all post-cells satisfying the connection type C, normalized by the number of realized (*b*_*ij*_ = 1) connection under C,

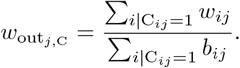

Similarly, the average realized summed monosynaptic inbound PSD for a post-cell *i* is given by

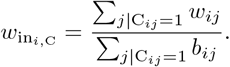

#### 5.11.2 Disynaptic Metrics

In examining the inhibition in sets of cells that share assembly membership, and in sets that do not, we were primarily interested in describing inhibition driven by the excitatory activity of an assembly’s member cells. This moved us from the realm of monosynaptic connection analysis into an analysis of chains. Many aspects of our definition remained unaltered. *A* remained the set of all assemblies, with *A*_*j*_ the subset of assemblies that included pre-cell *j*, and *A*_*i*_ the subset of assemblies that included post-cell *i*. The binary indicators indicating the assembly relationship between cells *i* and *j* remained unaltered.

But rather than simply using the monosynaptic weight between neurons *j* and *i* where *j* ≠ *i*, we defined *w*_*ikj*_ as the product of the pairwise summed post-synaptic densities *w* in a three-cell chain motif with *j* as the first cell, an interneuron *k* as the second cell, and *i* as the third cell. Thus,

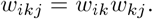

And similar to our monosynaptic analysis, *b*_*ikj*_ was an indicator variable that took the value 1 if at least one disynaptic chain existed between *j, k*, and *i* and 0 otherwise.

Building on this definition, we outlined the method of normalization and metrics for disynaptic chain analysis, which accounted for both the intermediate and final (or first) cells in the chain. In this analysis, each pre-cell and post-cell was a coregistered excitatory neuron with an extended axon, consistent with the monosynaptic analysis. The intermediary cell in disynaptic chains, however, did not need to be coregistered or possess an extended axon.

Let *n*_*e*_ represent the number of excitatory cells and *n*_*i*_ represent the number of inhibitory cells from the set of all cells in the all-all connectome. Define |*k*| be the number of potential middle partners. For inhibitory chains, |*k*| = *n*_*e*_ as the middle cell is inhibitory. For excitatory chains, the middle cell cannot be the first or final cell, so |*k*| = *n*_*e*_ − 2.

The probability of disynaptic outbound connection for a pre-cell *j* was calculated as the proportion of realized disynaptic connections under a given connection type C ∈ {Shared, Disjoint}, normalized by the total number of potential chains satisfying the connection type,

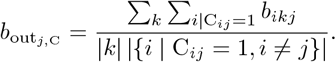

Similarly, the probability of disynaptic inbound connection for a post-cell *i* was defined as

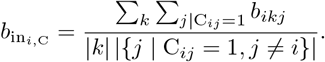

For nonzero strength of connection, we computed the average realized summed disynaptic outbound PSD for a pre-cell *j* as the total PSD across all chains satisfying the connection type C, normalized by the number of realized (*b*_*ikj*_ = 1) chains under C,

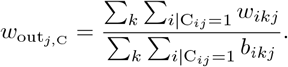

Similarly, the average realized summed disynaptic inbound PSD for a post-cell *i* was given by

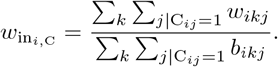

To facilitate statistical testing, we defined collections of metrics based on connection type C for both monosynaptic and disynaptic analyses. For disynaptic sets, the indices *ij* are replaced with *ijk* appropriately.

#### 5.11.3 Set Definitions

The following sets were defined to evaluate connectivity metrics:

- The set of nonzero pairwise connection strengths under a given connection type C,

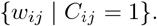
- The set of outbound probabilities of connection for each pre-cell under a given connection type C,

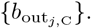

Similar collections were defined for inbound probability of connection, and for inbound and outbound nonzero average connection strengths by replacing the metric accordingly.
- For inbound or outbound metrics, paired sets were constructed by including only post- or pre-cells with at least one fulfilled connection under both the Shared and Disjoint connection types. Let

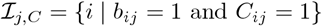

be the set of post-cells connected to pre-cell *j* under connection type *C*. The set of pre-cells included in the paired comparison was then defined as:

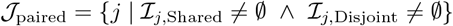

and the paired set of outbound metrics was:

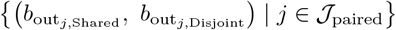

Similar paired collections were constructed for inbound probabilities and both inbound and outbound nonzero average connection strengths, as well as for metrics where pre- or post-cells appear in both the Assembly and the Non-Assembly groups.

These sets provided the basis for the statistical tests used to compare metrics across connection types.

#### 5.11.4 Statistical Tests

We performed one-way statistical tests at *α* = 0.05 to compare the Shared and Disjoint groups, as well as to compare the Assembly and Non-Assembly groups. All alternative hypotheses predict Shared *>* Disjoint or Assembly *>* Non-Assembly, except for tests involving di-synaptic Inhibitory chain sets, in which the alternative hypotheses predict Shared *<* Disjoint or Assembly *<* Non-Assembly. We ran the following tests:

- For unpaired sets, we use a one-sided Wilcoxon Rank-Sum test
- For paired sets, we use a one-sided paired Wilcoxon Signed-Rank test to compare metrics within cells appearing in both groups.

For pairwise binary connectivity, we created a contingency table to compare the frequencies of successful and failed connections across connection types:

| Connection Type | Successful Connections ($b_{ij} = 1$) | Failed Connections ($b_{ij} = 0$) |
| --- | --- | --- |
| Shared | $\sum_{ij\vert \text{Shared}_{ij}=1} b_{ij}$ | $\sum_{ij\vert \text{Shared}_{ij}=1} (1 - b_{ij})$ |
| Disjoint | $\sum_{ij\vert \text{Disjoint}_{ij}=1} b_{ij}$ | $\sum_{ij\vert \text{Disjoint}_{ij}=1} (1 - b_{ij})$ |

Then, we performed a Chi-Squared Test of Independence at *α* = 0.05 to determine if pairwise connection frequency differs across connection types.

Finally, to examine the functional correlates of the di-synaptic inhibitory chain findings, we calculated the Pearson’s correlation coefficient between the summed feed-forward inhibitory weights of disjoint cells connecting an assembly pair (see Methods 5.11.2) and the correlation scores between the assembly pair’s coactivity traces (see Methods 5.8), and examined significance.

#### 5.11.5 Tail Analysis

In addition to the mono-synaptic and di-synaptic analyses, we performed a “tail” analysis to investigate whether the proportion of Shared versus Disjoint connections differs between all pairwise connections and those classified as “tail” connections.

To identify “tail” connections, we modeled the distribution of connection strengths using a Gaussian Mixture Model (GMM) with *k* = 2 components. The model was initialized via k-means clustering to estimate the weights, means, and standard deviations of each component. The decision boundary separating the two Gaussian components was calculated as the intersection of their weighted probability density functions, derived using a quadratic equation based on the GMM parameters. Connections with values greater than or equal to the decision boundary were classified as “tail” connections. We present the model fit and evaluation as well as the tail boundary in the supplemental figure section (see Supp. Fig. 8).

Once the tail connections were identified, we compared the proportions of Shared and Disjoint connections in this subset to their proportions in the full dataset using a Chi-Squared Goodness-of-Fit Test at *α* = 0.05. This test considered only the Shared and Disjoint groups, with expected proportions calculated relative to the total counts of these two groups in the full dataset.

#### 5.11.6 Centrality analysis

Centrality analysis was used to quantify whether in a given network, assembly cells were more likely to be central to the network than non-assembly cells. This analysis gave further insights into the role of assembly cells in higher-order connectivity. To do this, we measured different centrality metrics for assembly and non-assembly cells, namely, indegree centrality, outdegree centrality, closeness centrality, and betweenness centrality.

In a graph, the centrality of a node refers to its tendency to connect and generally influence other nodes within the network [76]. We developed a directed graph *G* = (*N, E*), using the binary connectome such that |*N*| represents the number of cells in the connectome and *E* represents the binary, directed connections between all cell pairs. The total number of outbound synaptic connections are given by ∑ _*n*_ deg^+^(*n*) and inbound synaptic connections are given by ∑ _*n*_ deg^−^ (*n*). Normalizing these connections, indegree centrality was calculated as

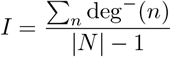

and outdegree centrality was calculated as

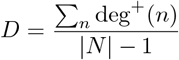

where *n* ∈ *N*.

A monosynaptic connection, *e* = (*j, i*) ∈ *E* between a pre-cell *j* and post-cell *i*, will have *i* as the head and *j* as the tail end of the connection, where *j* and *i* ∈ *N*. The path between a pre-cell *j* acting as a source neuron and a post-cell *i* acting as a target neuron, is the alternating sequence of cells and connections starting from *j* and ending at *i*, with each cell before a connection in the sequence being a pre-cell and each cell after a connection being a post-cell. The number of monosynaptic connections within the path indicates the length of the path. The shortest path between cells *j* and *i* is the minimum length between the two cells. The shortest path, *ξ*_*ij*_, is also referred to as the geodesic path.

Based on this, the closeness centrality for a given cell *i* was calculated as the reciprocal of the sum of the shortest paths, or distances between the post-cell *i* and all other pre-cells *j* in the graph,

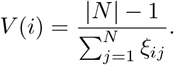

For a given cell *k*, the betweenness centrality [77] was calculated as

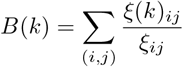

where *ξ*(*k*)_*ij*_ is the number of shortest paths between pre-cell *j* and post-cell *i* that pass through cell *k*. This value was also normalized to fall between 0 and 1.

## 6 Code Availability

All code and data are available at this GitHub link https://github.com/AllenInstitute/HebbsVision.

It should be noted that, for all code of our own generation, a fixed random seed is used for reproducibility. Libraries used for analysis and plotting of results include: numpy [78], pandas [79], scikit-learn [80], upsetplot [33], networkx [81], dotmotif [82], matplotlib [83], seaborn [84], and Raincloud [85].

## 7 Acknowledgements

We wish to thank the Allen Institute for Brain Science founder, Paul G. Allen, for his vision, encouragement, and support. S.B. has been supported by NIH R01EB029813. S.M. has been in part supported by NSF 2223725, NIH R01EB029813, and RF1DA055669 grants.

## 8 List of Contributions

| Name | Contributions |
| --- | --- |
| J Wagner-Carena | Conception and Design, Functional and Structural Data Analysis, Data Interpretation, Writing, Revision |
| S Kate | Data Interpretation, Chain and Graph Theoretic Data Analysis, Writing, Revision |
| T Riordan | Data Interpretation, Statistical Data Analysis, Writing, Revision |
| R Abbasi-Asl | Data processing, Revision, Writing |
| J Aman | Coregistration (Data Analysis), Writing |
| A Amster | Functional ROI extraction, team PIKA |
| AL Bodor | Data Acquisition: EM Histology/Sectioning; Revision |
| D Brittain | Data Acquisition: EM Imaging; Revision |
| JA Buchanan | Data Acquisition: EM Histology; Revision |
| MA Buice | Data processing: ROI extraction; Revision |
| DJ Bumbarger | Data Acquisition: EM Sectioning; Revision |
| F Collman | Interpretation, Dataset Resources and Infrastructure; Revision |
| NM da Costa | Data Processing; Interpretation; Writing, Revision |
| DJ Denman | Data Acquisition: Imaging, stimuli |
| SEJ deVries | Scientific leadership, optical physiology |
| E Joyce | Data Processing: Skeleton generation, layer definition, soma&nucleus feature extraction; Revision |
| D Kapner | Data Processing: EM alignment; Revision |
| CW King | Software development: allen_v1dd library |
| JD Larkin | Data Acquisition: Functional Imaging |
| D Liu | Software development: allen_v1dd library |
| G Mahalingam | Data Processing: EM Image Processing; Revision |
| D Millman | Stimulus Creation, Data Analysis |
| J Mölter | SGC Algorithm, Writing, Revision |
| C Morrison | Functional ROI extraction, team PIKA |
| RC Reid | Interpretation; Dataset Resources and Infrastructure, Revision |
| CM Schneider-Mizell | Interpretation; Dataset Resources and Infrastructure, Revision |
| D Scott | Functional ROI extraction, team PIKA |
| S Suckow | Project Management |
| KT Takasaki | Data Acquisition: Functional Imaging |
| M Takeno | Data Acquisition: EM Histology/Sectioning; Data Processing: coRegistration; Revision |
| R Torres | Data Processing: EM Image Processing/Coregistration; Revision; Writing |
| D Vumbaco | Proofreading Project Management |
| J Waters | Microscopy team lead |
| DG Wyrick | Software development: allen_v1dd library |
| W Yin | Data Acquisition: EM Imaging; Revision |
| J Zhuang | Data processing |
| S Mihalas | Conception and Design, Data Interpretation, Writing, Revision |
| S Berteau | Conception and Design, Data Analysis, Data Interpretation, Writing, Revision |

## 9 Statistical Tests

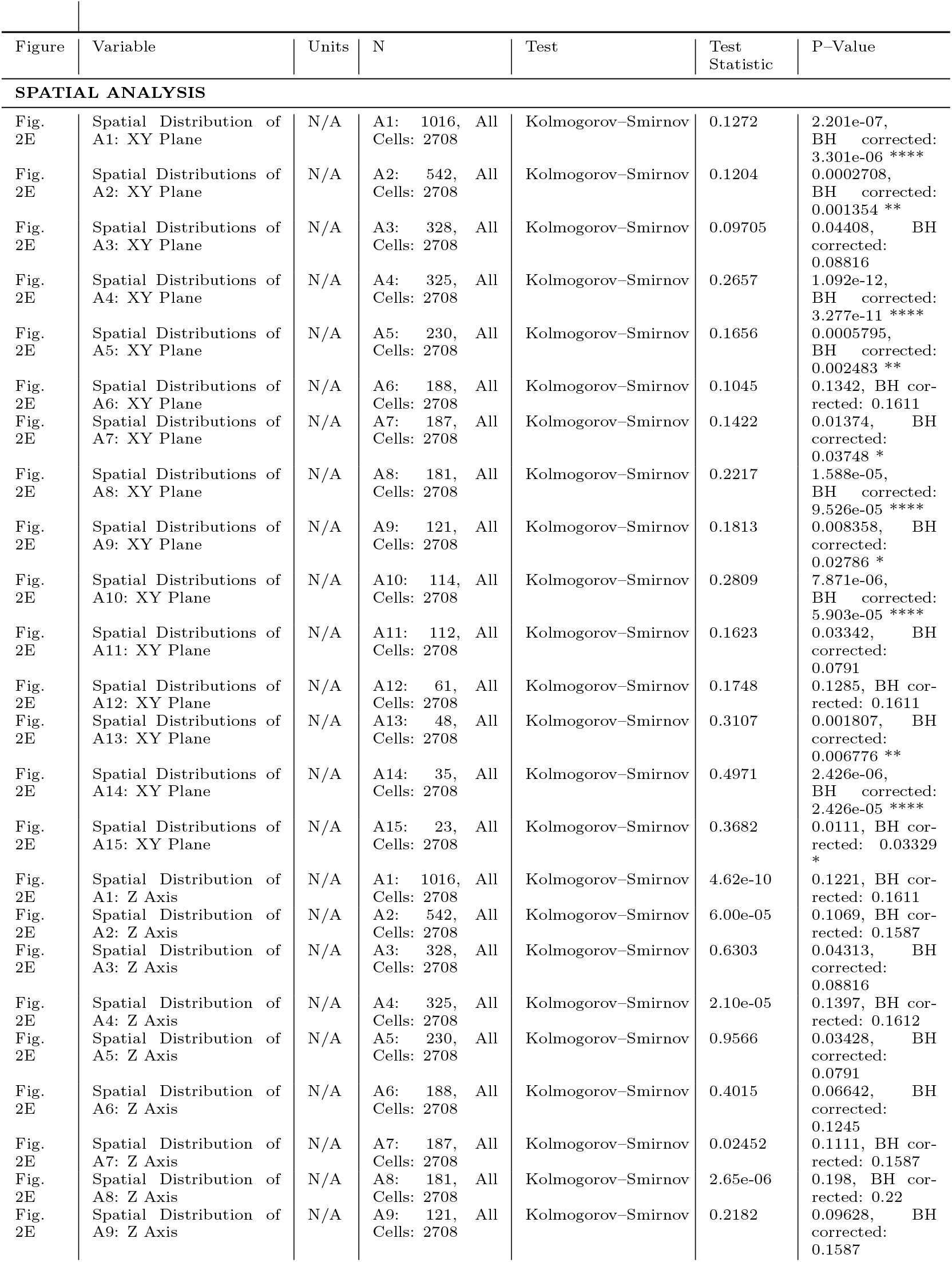
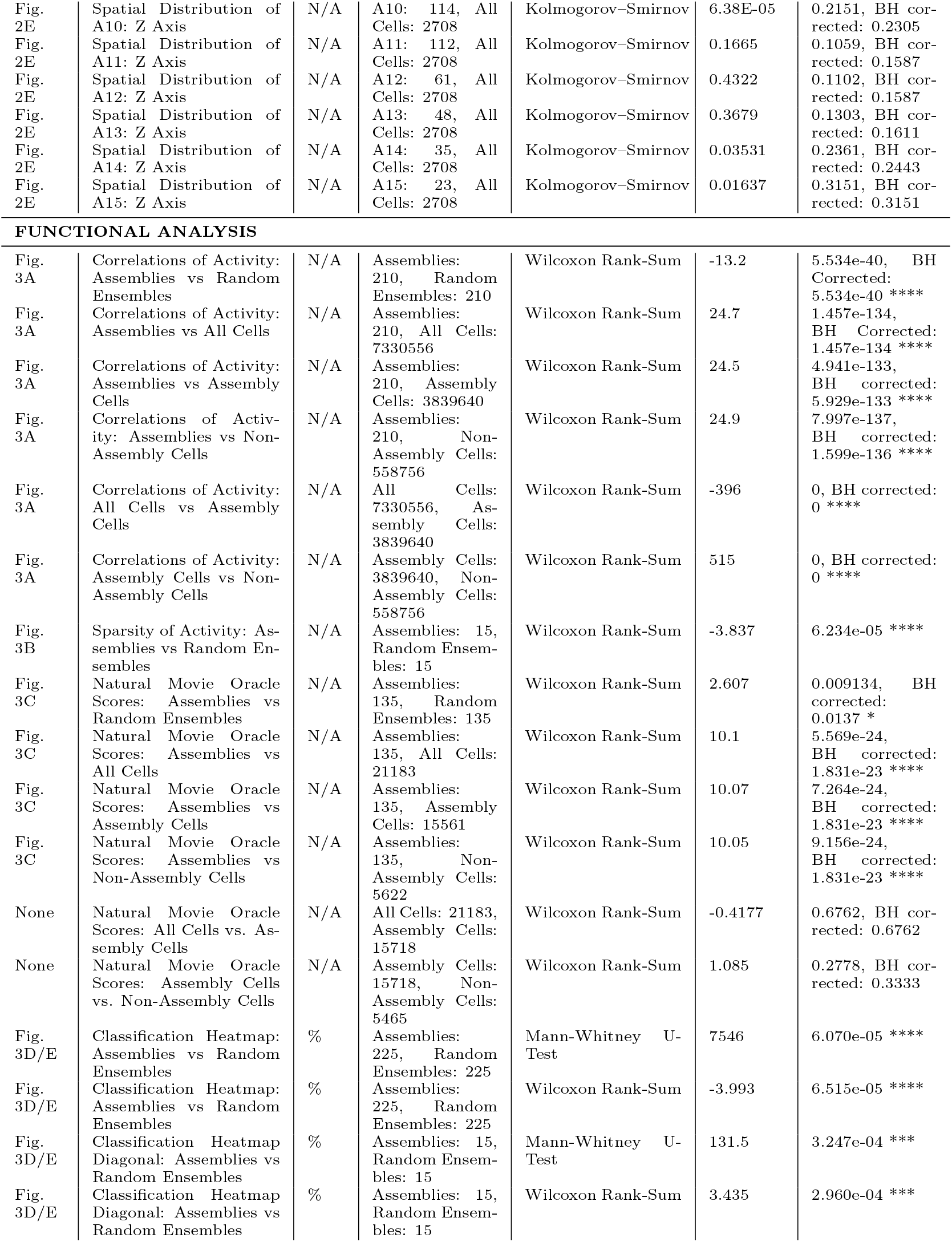
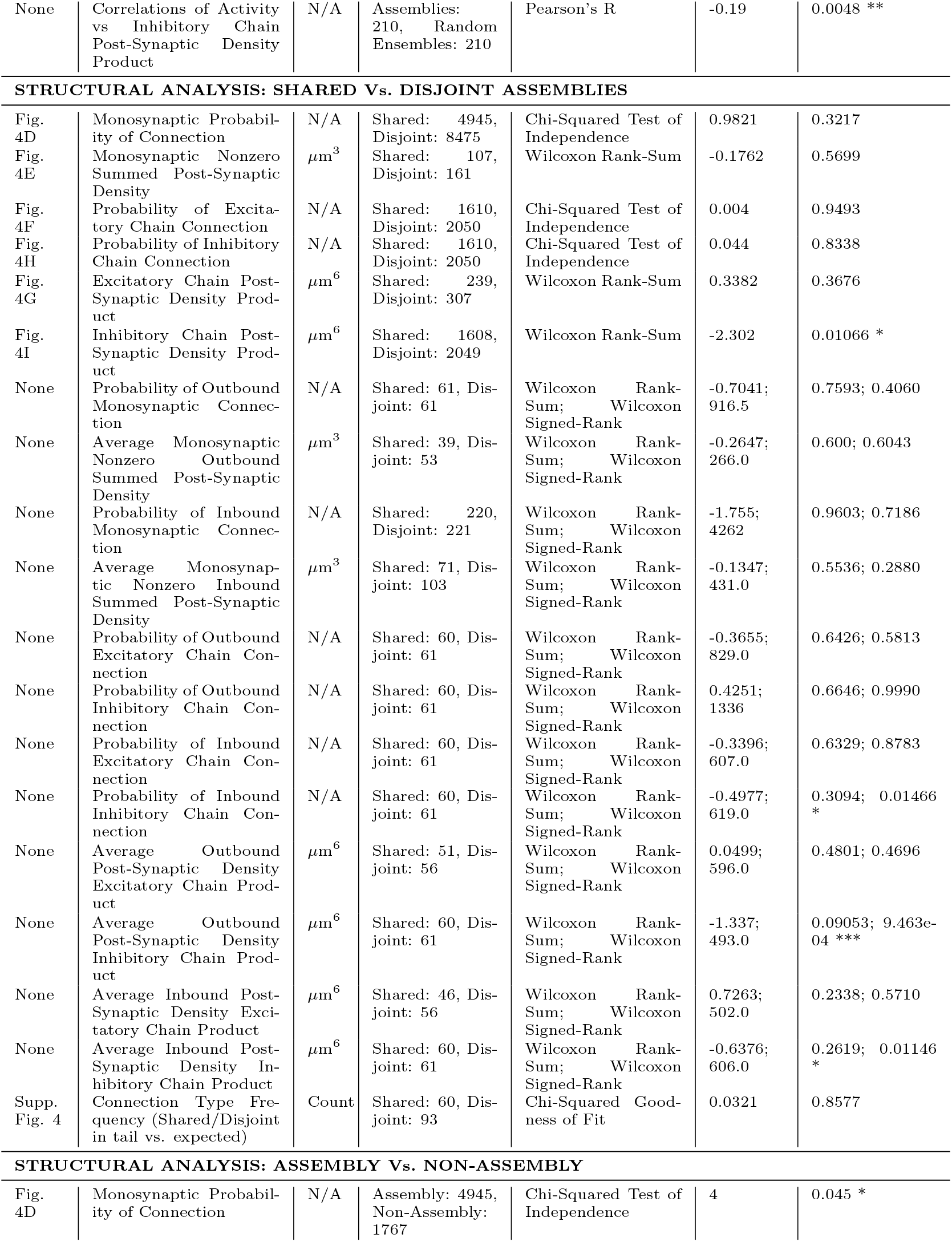
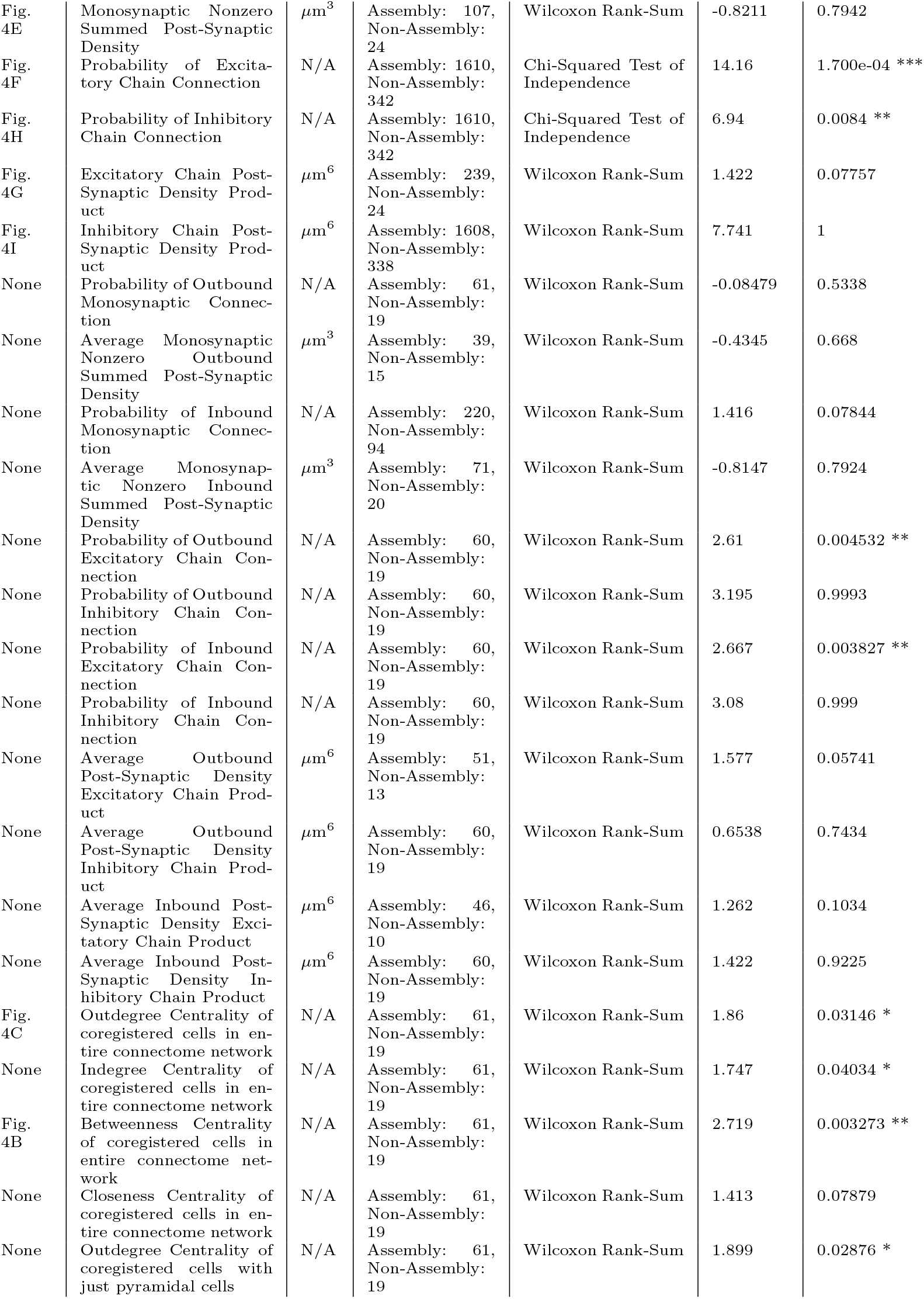
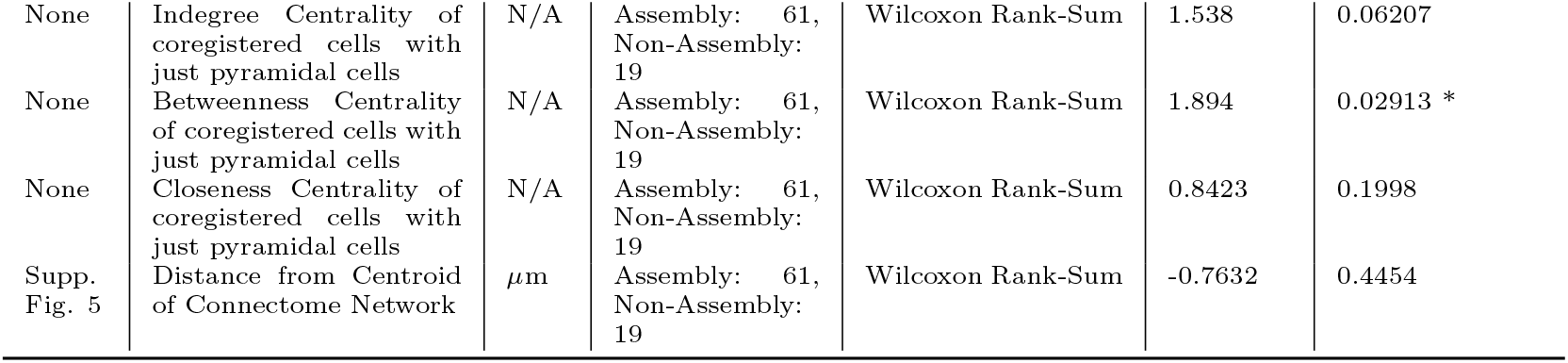

## 10 Extended Data

**Supplementary Figure 1.**
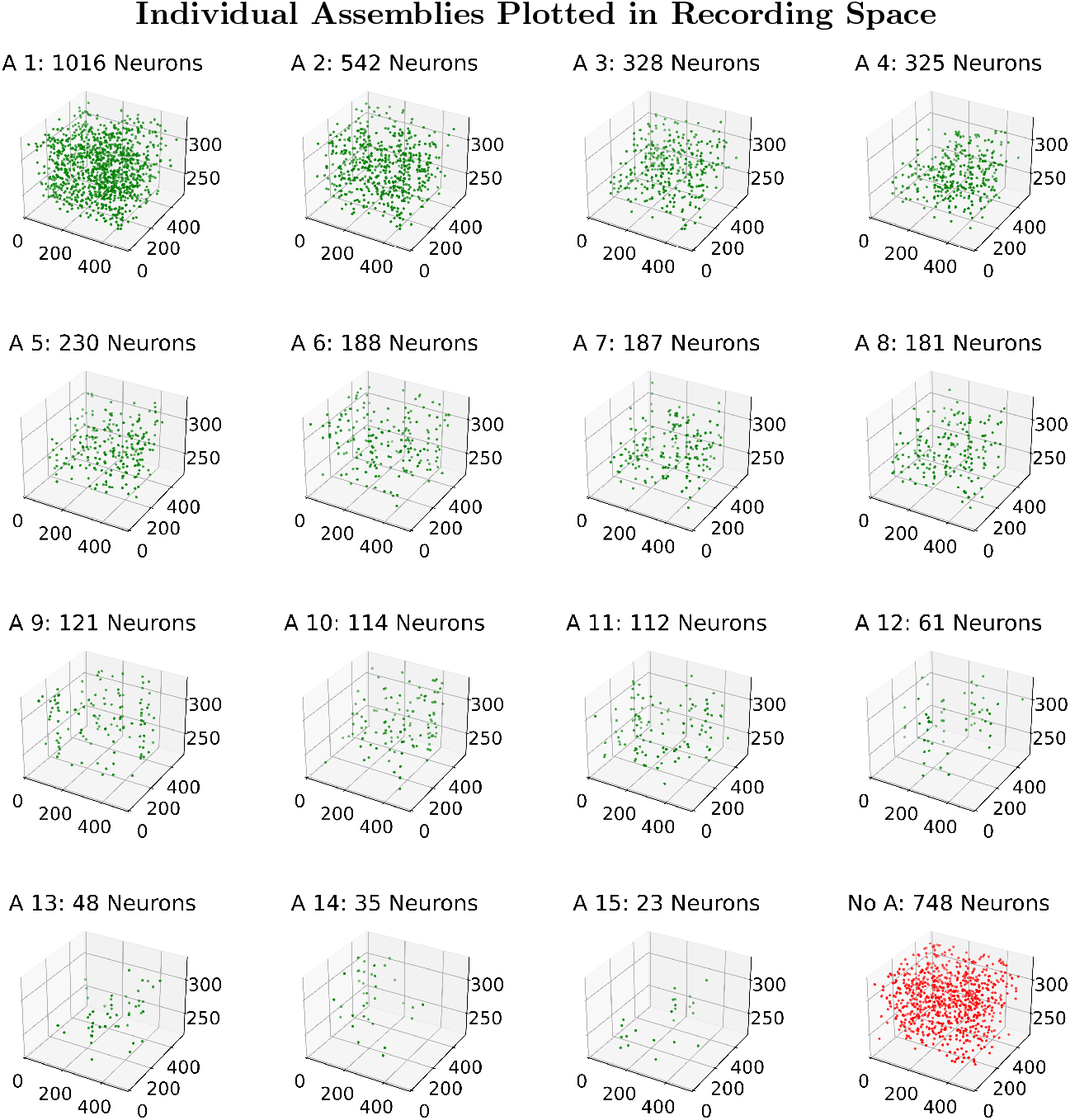
Spatial positions of individual extracted assemblies in the three-dimensional recording field. Every subplot provides an isolated view of an assembly, ordered by size, visualized in the optical imaging recording space. Each individual point refers to an identified excitatory neuron. The plot of neurons assigned to no assemblies, ‘No A’, is also shown in red (bottom-right). Sub-plot axes refer to the three spatial dimensions of the recording field, with units in micrometers. Sub-plot titles also include the size of each assembly. In total, 1960 neurons were assigned to assemblies, and 748 were not. Assemblies are typically spatially spread out throughout the recording field.

**Supplementary Figure 2.**
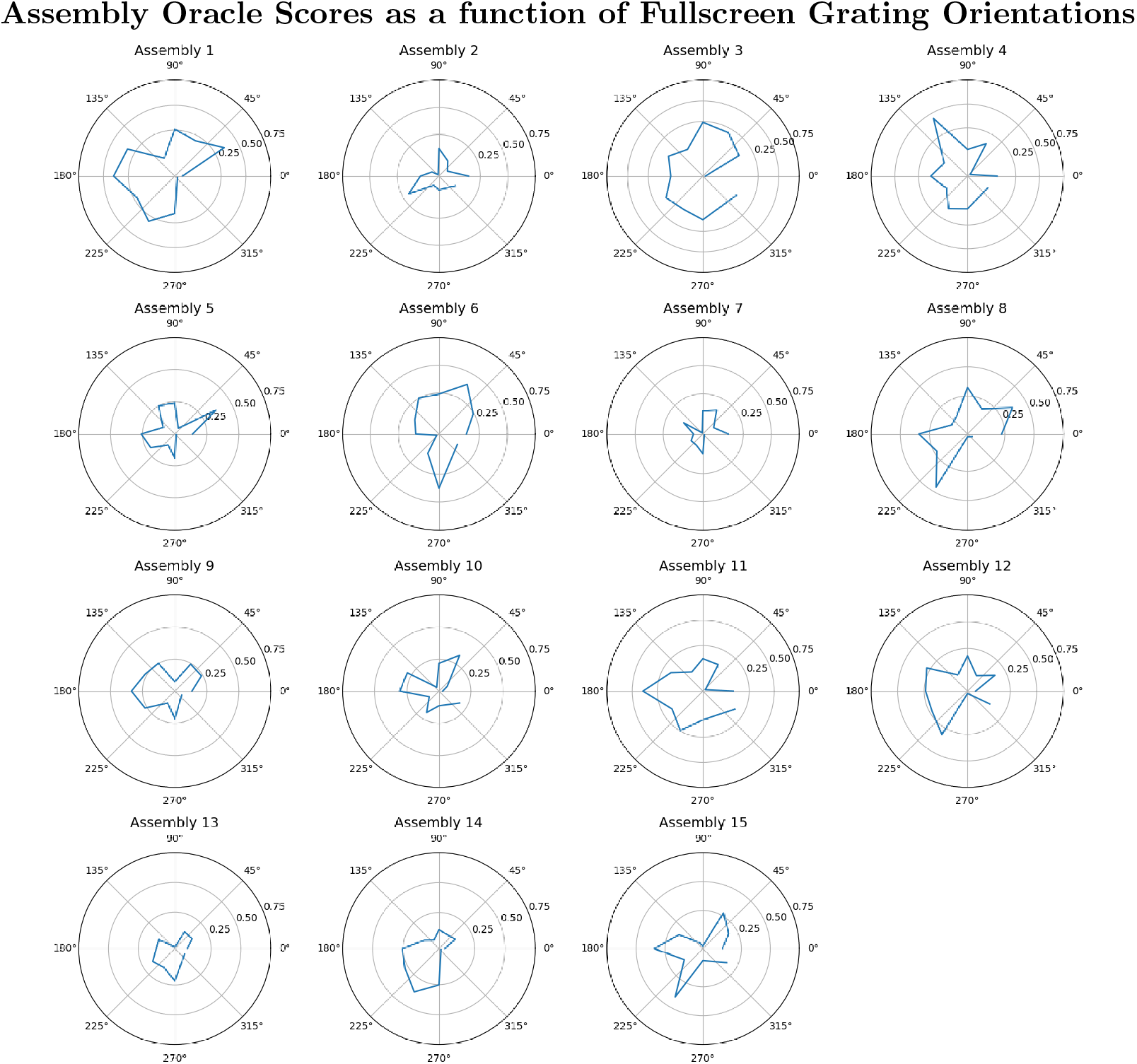
Individual Assemblies Oracle Scores to Fullscreen Gratings. Every sub-plot provides an isolated visualization of the reliability in an assembly’s response with respect to the orientation of fullscreen gratings. Orientation of gratings is represented by a polar plot. Reliability is measured through the Oracle score metric (see Methods 5.3). The scores of assembly coactivity trace in response to gratings are typically lower than those seen in natural movies (Fig. 3C), but the results are still indicative of tuning properties in these functional populations. Notably, some assembly traces seem to be highly reliable to particular orientations, similar to the orientation receptive fields of simple cells in the primary visual cortex.

**Supplementary Figure 3.**
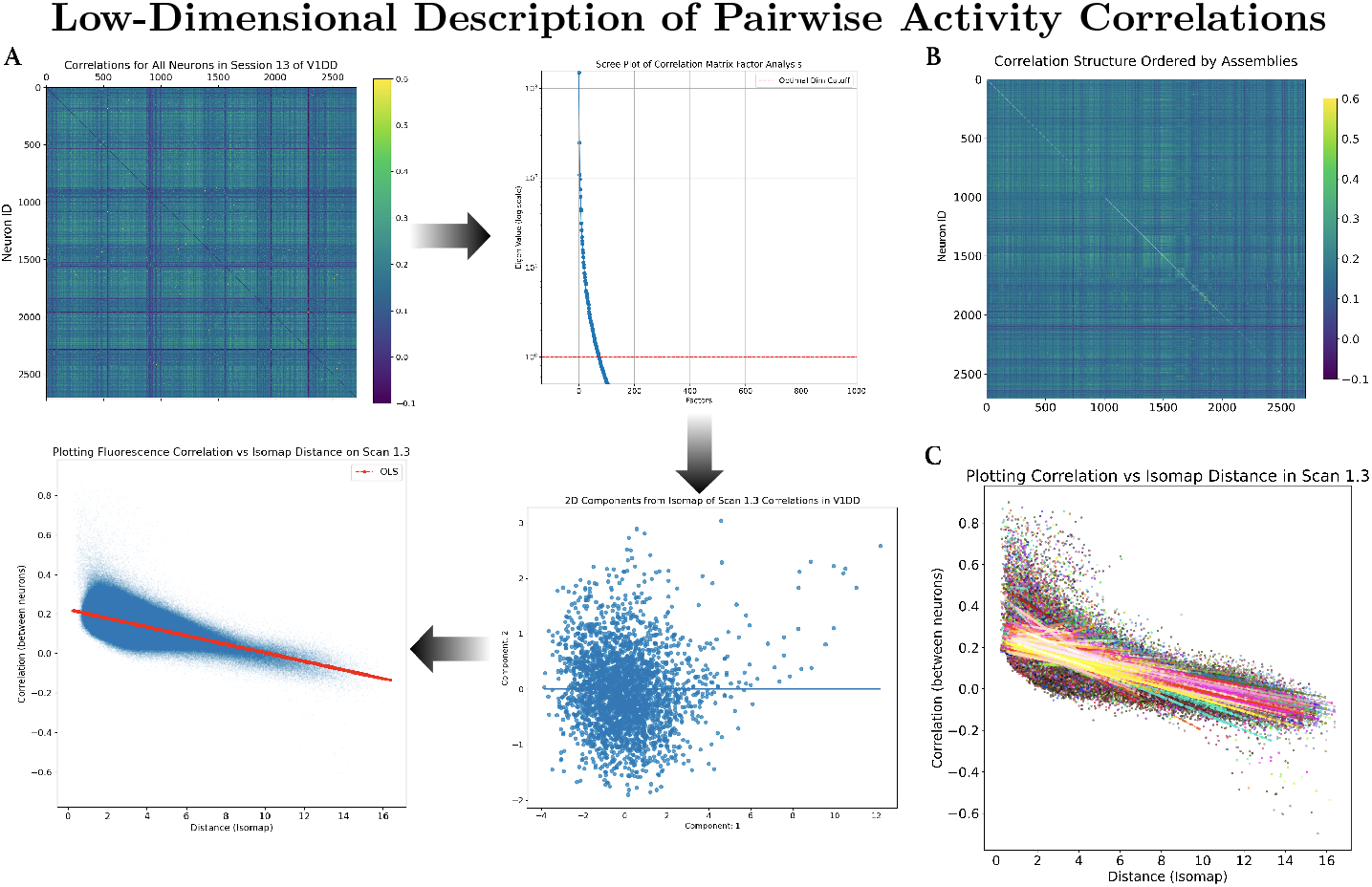
Fitting Regressions to establish low-dimensional descriptions of pairwise activity correlations of cells. This framework was used to choose the optimal hyperparameters from Scan Volume 4, with those parameters later being chosen for Scan Volume 3. **(A)**. Pipeline to produce our regression fits to activity correlations. *Top-left* : The procedure begins with the pairwise correlation matrix of the whole scan throughout the hour recording. *Top-right* : Factor analysis reveals which dimensions of the original data matrix explain the most variance. This is done by plotting the eigenvalue of each factor, or dimension. The dotted red line is a horizontal marker of the eigenvalue of one, a threshold for considered dimensions justified by the Kaiser criterion. We ensure that at least ninety percent of the total variance is left explained in the final embedding. Less than fifty factors, or dimensions, are needed to do so. *Bottom-right* : With these dimensions in mind, we apply Isomap to produce a non-linear low-dimensional manifold of the original correlation matrix. This plot illustrates the first and second components from that embedding. *Bottom-right* : Finally, our lowdimensional description of the activity correlation, or our activity correlation space, is produced by plotting the distance of neurons across the Isomap embedding with respect to their pairwise correlations. The OLS fit (solid-red line) is a statistical model exemplifying how well the system continuum is able to describe this activity correlation space. **(B)** Correlation matrix of the whole scan with neurons ordered by assemblies. Cells on each axis were ordered by their largest corresponding assembly, from ‘A 1’ to ‘A 15’. **(C)** OLS fits were developed that corresponded to assembly assignments with pairwise cells. This implies that multiple OLS fits were performed to account for all assignments. Regression fits to different assembly assignments are compared through the Akaike information criterion.

**Supplementary Figure 4.**
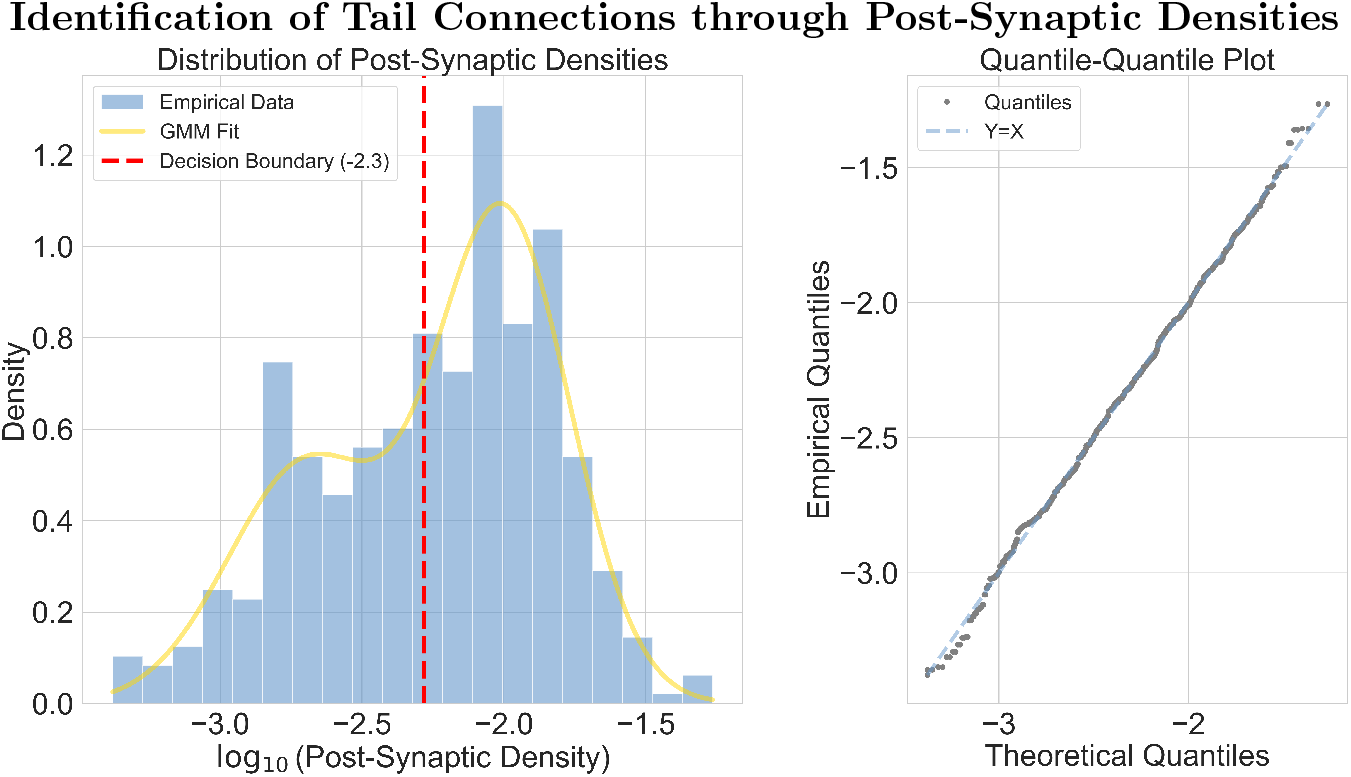
Histogram of connection strengths (log_10_-scaled) with a Gaussian Mixture Model fit overlaid (see Methods 5.11.5), and a Quantile-Quantile plot evaluating the fit. The red dashed line indicates the decision boundary separating the two Gaussian components, used to classify “tail” connections.

**Supplementary Figure 5.**
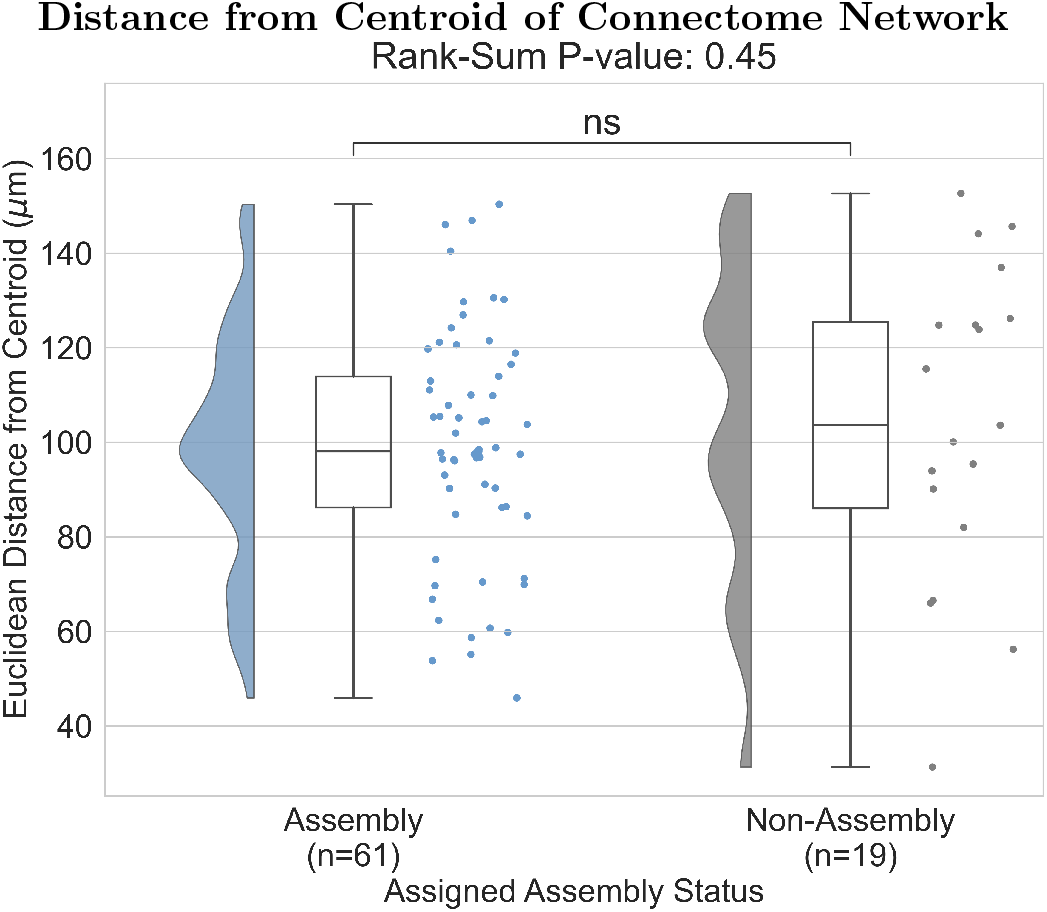
A raincloud plot comparing the distance from the centroid of the connectome network (Fig. 4A), demonstrating no difference between the distribution of distances with neurons assigned to an assembly and the distribution of those assigned to no assembly (Wilcoxon Rank-Sum p-value: 0.45).

